# Genetic variation in *CSMD1* affects amygdala connectivity and prosocial behavior

**DOI:** 10.1101/2020.09.27.315622

**Authors:** KC Bickart, V Napolioni, RR Khan, Y Kim, A Altmann, J Richiardi, M Newsom, S Sadaghiani, T Banaschewski, ALW Bokde, EB Quinlan, S Desrivières, H Flor, H Garavan, P Gowland, A Heinz, B Ittermann, J-L Martinot, M-L Paillère Martinot, E Artiges, F Nees, D Papadopoulos Orfanos, T Paus, L Poustka, JH Fröhner, MN Smolka, H Walter, R Whelan, G Schumann, B Ng, MD Greicius, IMAGEN Consortium

**Author notes:** Correspondence can be directed to: Kevin Bickart, 780 Welch Road, Palo Alto, Ca 94304, e. Contributed equally (KC Bickart, V Napolioni).

## Abstract

The amygdala is one of the most widely connected structures in the primate brain and plays a key role in social and emotional behavior. Here, we present the first genome-wide association study (GWAS) of whole-brain resting-state amygdala networks to discern whether connectivity in these networks could serve as an endophenotype for social behavior. Leveraging published resting-state amygdala networks as *a priori* endophenotypes in a GWAS meta-analysis of two adolescent cohorts, we identified a common polymorphism on chr.8p23.2 (rs10105357 A/G, MAF (G)=0.35) associated with stronger connectivity in the medial amygdala network (beta=0.20, *p*=2.97×10^−8^). This network contains regions that support reward processes and affiliative behavior. People carrying two copies of the minor allele for rs10105357 participate in more prosocial behaviors (t=2.644, *p*=0.008) and have higher *CSMD1* expression in the temporal cortex (t=3.281, p=0.002) than people with one or no copy of the allele. In post-mortem brains across the lifespan, we found that *CSMD1* expression is relatively high in the amygdala (2.79 fold higher than white matter, *p*=1.80×10^−29^), particularly so for nuclei in the medial amygdala, reaching a maximum in later stages of development. Amygdala network endophenotyping has the potential to accelerate genetic discovery in disorders of social function, such as autism, in which *CSMD1* may serve as a diagnostic and therapeutic target.

---

The amygdala is one of the most widely-connected structures in the primate brain. It is a hub for socioemotional information with connections to a diverse array of cortical and subcortical structures involved in perceiving social cues, affiliating with friends, and avoiding foes (Bickart, Dickerson, & Barrett, 2014). Previous work using a novel approach to resting-state fMRI revealed that the amygdala’s broad array of connections can be parsed into three large-scale brain networks (Bickart, Hollenbeck, Barrett, & Dickerson, 2012) that support three distinct, fundamental domains of socioemotional behavior (Bickart, Brickhouse, et al., 2014). Elucidating the genetic architecture of these networks is a necessary step in translating this research to the clinical care of patient populations marked by amygdala dysfunction and associated neurobehavioral disruptions.

The amygdala’s structural and functional properties have become a major focus in imaging genetics, investigated as endophenotypes to elucidate genetic variance in systems relevant to the amygdala’s role in socioemotional behavior (Buckholtz et al., 2007; Canli & Lesch, 2007; Meyer-Lindenberg et al., 2006a). Despite this progress, genome-wide studies of amygdala circuitry have not been performed, and candidate gene-based studies have focused on specific connections between the amygdala and a limited number of connected regions (Buckholtz et al., 2008; Kruschwitz et al., 2015), none yet exploring a more comprehensive model of amygdala circuitry. Here, we performed a genome-wide association study (GWAS) on the three previously published large-scale resting-state networks of the amygdala (Figure 1) to discern whether connectivity strength in these networks can serve as endophenotypes for normal variation in socioemotional behavior. The endophenotype, or intermediate phenotype, concept posits that biomarkers may be more likely than behavioral phenotypes to reveal novel genetic variants relevant to psychiatric disorders (Meyer-Lindenberg & Weinberger, 2006). Prior imaging genetic studies have used this approach to make discoveries in a variety of disorders, such as autism (Fakhoury, 2018) and schizophrenia (Li et al., 2017).

**Figure 1.**
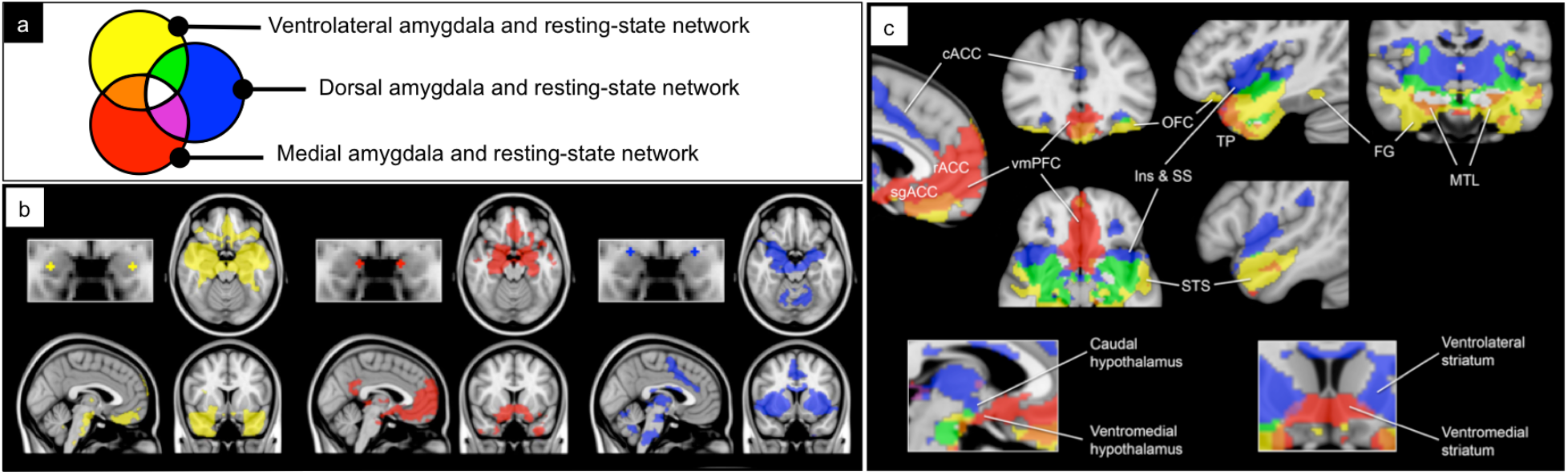
Resting-state seeds and networks of the amygdala. Previously published (Bickart et al., 2012) one sample group mean significance maps for each amygdala seed color-coded by seed (a) displayed in standard views (N=89) for each seed separately (b) and all three seeds together (c) demonstrating unique and overlapping connectivity patterns. The maps are binarized at *p*<10^−5^ and overlaid on a T1 MNI152 0.5mm template brain in radiologic convention to demonstrate the distinct and shared connectivity across maps. Abbreviations: lOFC, lateral orbitofrontal cortex; vmPFC, ventromedial prefrontal cortex; cACC, caudal anterior cingulate cortex; Ins, insula; SS, somatosensory operculum; STS, superior temporal sulcus; dTP, dorsal temporal pole; OFC, orbitofrontal cortex; rACC, rostral anterior cingulate cortex; sgACC, subgenual anterior cingulate cortex; MTL, medial temporal lobe; FG, fusiform gyrus; vTP, ventral temporal pole; vlSt, ventrolateral striatum; vmSt, ventromedial striatum.

Our *a priori* amygdala network endophenotypes (Figure 1) emanate from nuclei within the medial amygdala (red), dorsal amygdala (blue), and ventrolateral amygdala (yellow), as defined previously using an anatomically-driven approach to resting-state fMRI (Bickart et al., 2012). The medial amygdala network contains reward-related regions within the ventromedial prefrontal cortex, nucleus accumbens, ventromedial hypothalamus, ventral tegmental area and medial temporal lobe implicated in guiding goal-directed behavior in general and affiliation with others in the social domain (red in Figure 1). The dorsal amygdala network contains nociceptive areas within the dorsal anterior cingulate cortex, insula, somatosensory operculum, ventral basal ganglia, hypothalamus, thalamus, and medial temporal lobe implicated in avoiding aversive stimuli in general and untrustworthy or threatening people in the social realm (blue in Figure 1). The ventrolateral amygdala network contains multimodal sensory association, affective, and memory areas of the temporal and orbitofrontal lobes implicated in the perception of affective sensory information in general and signals of communication in the social realm, such as faces, facial expressions, and body gestures (yellow in Figure 1).

## Methods

### Participants

We acquired data for our GWAS and endophenotype analyses from the I IMAGEN (G. Schumann et al., 2010) and Neurodevelopmental Cohort (PNC) (Satterthwaite et al., 2014) cohorts. For the GWAS meta-analysis, we specifically selected subjects that identified as Caucasian, had whole genome data, and had resting-state fMRI data (Table 1). For the endophenotype analysis, we selected independent samples of subjects within the cohorts that identified as Caucasian, had whole genome data with genotypes for our SNPs of interest, *but had not been included in the GWAS analysis* (Table 1). All other details regarding participant recruitment, exclusion/inclusion criteria, and consent can be found in the original publications for IMAGEN (G. Schumann et al., 2010) and PNC (Satterthwaite et al., 2014).

**Table 1.** Demographics and genotypes of subjects included in this study

|  | IMAGEN | PNC |
| --- | --- | --- |
| **Subjects included in the GWAS meta-analysis** |  |  |
| Mean age (SD) | 14 (0) | 14.6 (3.4) |
| Sample size (Female) | 315 (142) | 390 (189) |
| rs10105357 |  |  |
| A/A | 140 (60) | 148 (69) |
| A/G | 144 (68) | 198 (101) |
| G/G | 31 (14) | 41 (18) |
| rs35192025 |  |  |
| T/T | 130 (55) | 139 (64) |
| C/T | 144 (75) | 181 (95) |
| C/C | 38 (11) | 61 (26) |
| rs17671156 |  |  |
| G/G | 195 (86) | 245 (114) |
| G/T | 108 (53) | 130 (66) |
| T/T | 9 (2) | 12 (9) |
| **Subjects included in the endophenotype analysis** |  |  |
| Mean age (SD) | 14 (0) | 14 (4) |
| Sample size (Female) | 1358 (698) | 4427 (2195) |
| rs10105357 |  |  |
| A/A | 561 (284) | 1908 (934) |
| A/G | 650 (337) | 1894 (928) |
| G/G | 143 (76) | 466 (242) |
| rs35192025 |  |  |
| T/T | 557 (290) | 1539 (768) |
| C/T | 603 (308) | 2014 (1002) |
| C/C | 186 (93) | 683 (331) |
| rs17671156 |  |  |
| G/G | 856 (450) | 2728 (1359) |
| G/T | 436 (215) | 1394 (686) |
| T/T | 43 (22) | 171 (74) |

### Imaging acquisition

MR images for the IMAGEN cohort were acquired on 3T scanners from a variety of manufacturers, full details published previously (G. Schumann et al., 2010), with consistent parameters across sites. Resting-state fMRI parameters consisted of resting with eyes closed but not sleeping during a 6.5-minute gradient-echo, echoplanar sequence (GE-EPI: TR 2200ms, TE 30ms, flip angle 75°, matrix 64×64 voxels, 40 slices, 2.4mm slice thickness, 1mm slice gap, field of view 218×218mm, 3.4mm isotropic voxel size). Structural MRI parameters consisted of a T1-weighted sequence based on the well validated ADNI consortium protocols (http://adni.loni.usc.edu/methods/documents/mri-protocols/).

MR images for the PNC cohort were acquired on a 3T Siemens TIM Trio whole-body scanner, full details published previously (Satterthwaite et al., 2014). Resting-state fMRI parameters consisted of resting with eyes open and fixed on crosshair during a 6.2-minute GE-EPI sequence (TR 3000ms, TE 32ms, T1 1100ms, flip angle 90°, matrix 64×64 voxels, 46 slices, 3mm slice thickness, no slice gap, field of view 192×192mm, 3mm isotropic voxel size). Structural MRI parameters consisted of a T1-weighted MPRAGE sequence (TR 1810ms, TE 3.5ms, matrix 192×256, 160 slices, 1mm slice thickness).

### Imaging processing

For IMAGEN, we acquired preprocessed resting-state fMRI data from the consortium website, which already underwent slice-timing correction, motion correction, and spatial normalization to MNI space using SPM8 (http://www.fil.ion.ucl.ac.uk/spm/). For PNC, we acquired raw resting-state fMRI data and applied motion correction and spatial normalization to MNI space using ANTs (http://stnava.github.io/ANTs/). For both datasets, we then regressed out six linear head motion parameters, white matter and cerebrospinal fluid confounds, five principal components of high variance voxels derived using CompCor (Behzadi, Restom, Liau, & Liu, 2007), and their one-time sample shifted variants and subsequently performed bandpass filtering at 0.01-0.1Hz.

We computed connectivity in each of the three prior published amygdala networks (Bickart et al., 2012) for each subject. To do this, we computed a Pearson correlation between the BOLD signal time course averaged across all voxels belonging to the amygdala seed regions and the BOLD signal time course averaged across all voxels belonging to the amygdala networks (seeds and networks used can be seen in Figure 1). This produced three amygdala network correlation values per subject. We then standardized these values, setting the mean to 0 and variance to 1, and used them as the *a priori* endophenotypes in our GWAS meta-analysis.

### Genome-wide association meta-analysis

IMAGEN subjects were genotyped from blood samples on 610-Quad SNP and 660-Quad SNP arrays from Illumina (Illumina Inc., San Diego, CA). Most PNC subjects were genotyped from blood samples on the 550HH and 610-Quad SNP arrays from Illumina (Illumina Inc., San Diego, CA). Analyses were performed using PLINK 1.9 (Chang et al., 2015). GWAS datasets were available in different human genome builds (hg18 and hg19). Therefore, the hg18 datasets were lifted to hg19 version before QC processing using LiftOver chain available through the UCSC Genome Browser (Kent, 2002). Samples showing sex inconsistency and autosome missingness (> 5%) were excluded from the analysis.

Identity-by-descent (IBD) > 0.0625 cutoff, equivalent to 3^rd^ degree relative, was applied to exclude related subjects. Population structure outliers were determined by EIGENSOFT v.6.1 (Price et al., 2006) using pruned (r^2^ < 0.1) SNPs from each dataset. Before imputation, samples were processed using PLINK 1.9 (Chang et al., 2015) to remove SNPs with a MAF < 0.01, Hardy-Weinberg Equilibrium (HWE) P < 5 x 10^−6^ and missingness > 0.05. Each GWAS dataset was independently QCed, phased with SHAPEIT (Delaneau, Marchini, 1000 Genomes Project Consortium, & 1000 Genomes Project Consortium, 2014) and imputed with IMPUTE2 (Howie, Donnelly, & Marchini, 2009) considering the 1000 Genomes Project Phase 3 panel (1000 Genomes Project Consortium et al., 2015). After imputation, we excluded SNPs with a r^2^ quality score < 0.7, MAF < 0.05, and HWE (P < 5 x 10^−6^), yielding 4,825,426 SNPs shared across the two datasets.

Association testing was carried out using a logistic regression model implemented in PLINK 1.9 (Chang et al., 2015). The model was adjusted for subject’s age (for PNC only), sex, fMRI scanning site (for IMAGEN only) and the first three principal components from population structure analysis. Results from each dataset were fixed-effect meta-analyzed using GWAMA (Mägi & Morris, 2010) and the results were displayed using R libraries.

### Expression quantitative trait loci (eQTL) analysis

We performed eQTL analyses to test whether the top SNPs for each amygdala network affect gene expression in the brain for the gene in closest proximity. We used RNA-seq data from the temporal cortex of a subset of cognitively healthy subjects belonging to the Mayo RNA-Sequencing study (Allen et al., 2016) who had gene expression data and all covariates of interest (N=62). Covariates included age of death, sex, RNA integrity number, and post-mortem intervals (PMI). We retrieved data from the AMP-AD portal in the Sage Bionetworks Synapse project (synapse.org). For each of our top SNPs, we performed a linear regression to test the additive model as well as one-way ANOVA and post-hoc *t*-tests using the genetic model most representative of the distribution of the effect (e.g., recessive, dominant) controlling for covariates, accepting *p <* 0.05 as significant.

### Endophenotype analysis

We tested the effect of genotype for our top SNPs on behavioral phenotypes referable to each amygdala network in an independent subset of our cohorts. To do this, we acquired genotype and behavioral data for all Caucasian adolescents in PNC and IMAGEN that were not included in the GWAS (demographics by genotype detailed in Table 1). In each case, we performed a one-way ANOVA on the summary scores across levels of the variant allele followed by post hoc *t*-tests using the genetic model most representative of the distribution of the effect (e.g., recessive, dominant), accepting an alpha of 0.05 for significance. For significant findings, we repeated the *t*-tests on the unstandardized residual of each dependent variable after regressing out potential confounders (i.e., PNC: sex, age, and three genetic principal components; IMAGEN: sex, handedness, Tanner stage of puberty, imaging center, and three genetic principal components). Of note, tanner puberty stage was not available for the PNC cohort.

Given the function of the medial amygdala network, we hypothesized that carriers of the rs10105357-G polymorphism would demonstrate phenotypic differences in instruments assessing aspects of social affiliation. The IMAGEN cohort underwent assessment of such a phenotype, specifically, the “prosocial behavior” score generated from the Strengths and Difficulties Questionnaire (SDQ) (Goodman, 1997). It is a clinically validated (Silva, Osório, & Loureiro, 2015) composite of 5 items including “Considerate of other people’s feelings “, “Shares readily with other children (treats, toys, pencils, etc.)”, “Helpful if someone is hurt, upset or feeling ill”, “Kind to younger children”, and “Often volunteers to help others (parents, teachers, other children)”, rated as 0=not true, 1=somewhat true, 2=certainly true and summed for a total possible score of 10. We performed similar analyses of relevant phenotypes, bully behavior and face perception, for the top suggestive SNPs in the other two networks, as detailed in Supplementary Methods and Results.

### *CSMD1* expression in the brain and amygdala across the lifespan

We examined the spatial and temporal pattern of *CSMD1* expression in the brain using three datasets that are publicly available from the Allen Brain Sciences Institute. To assess the spatial pattern of *CSMD1’s* expression independent of age, we combined the adult and fetal microarray datasets from the Allen Brain Atlas (http://human.brain-map.org/) and the BrainSpan Atlas of the Developing Human Brain (http://www.brainspan.org/), respectively, to maximize the number of samples and degree of granularity in brain regions sampled. This allowed us to assess a range of brain regions throughout the cortex and subcortex (Figure 5a-b) and within subnuclei of the amygdala (Figure 5c-d) that make up the seed regions of our resting-state amygdala networks. These datasets were also normalized with similar methods and underwent quality control procedures, as described previously (Hawrylycz et al., 2012; Miller et al., 2014) and in technical white papers available at http://human.brain-map.org/ under the respective atlas’s “Documentation” tab. We downloaded the normalized z-scores for the adult brains at http://human.brain-map.org/, which originate from 3 probes, 6 brains spanning 24 to 57 years old, and 1202 tissue samples from left and right hemispheres making up 169 brain regions (Table 2). We downloaded the normalized z-scores for the fetal brains at http://www.brainspan.org/lcm/, which originate from the same 3 probes as the adult dataset, 4 brains spanning 15 to 21 weeks post-conception, and 1203 tissue samples from the left hemisphere making up 516 tissue regions (Table 2). We then pooled expression data across probes and tissue samples into common top level structure groups by cortical regions containing the following word in the structure name, “frontal”, “temporal”, “parietal”, “occipital”, “insular”, “cingulate”, “somatosensory”, and “cerebell” as well as subcortical regions making up the amygdala, hippocampal formation, basal forebrain, striatum, hypothalamus, brainstem, and white matter. As an exploration of anatomical specificity, we compared *CSMD1* expression between the amygdala and white matter and between the amygdala subregions using independent samples *t*-tests, accepting *p* < 0.05 as significant.

**Table 2.**
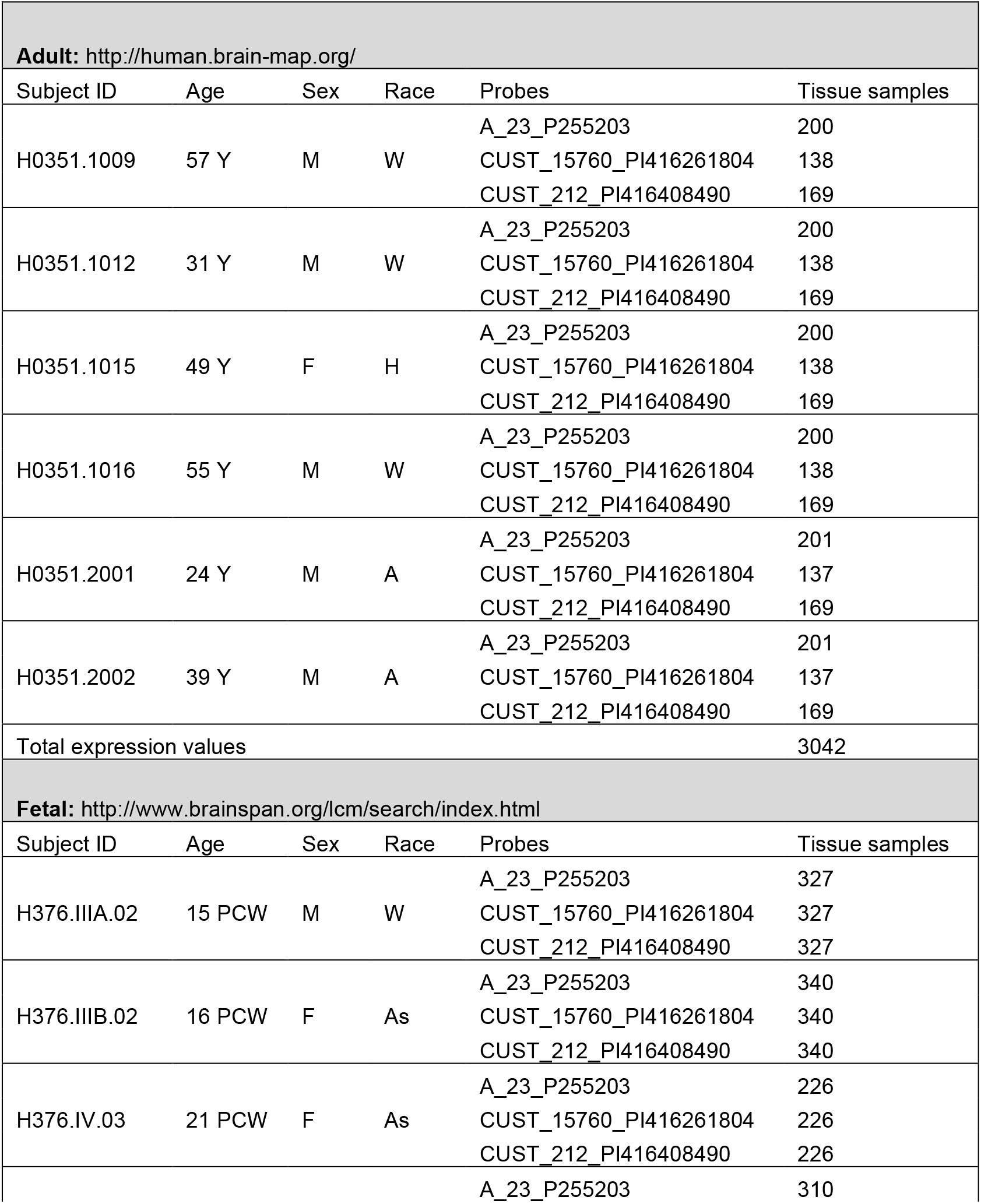
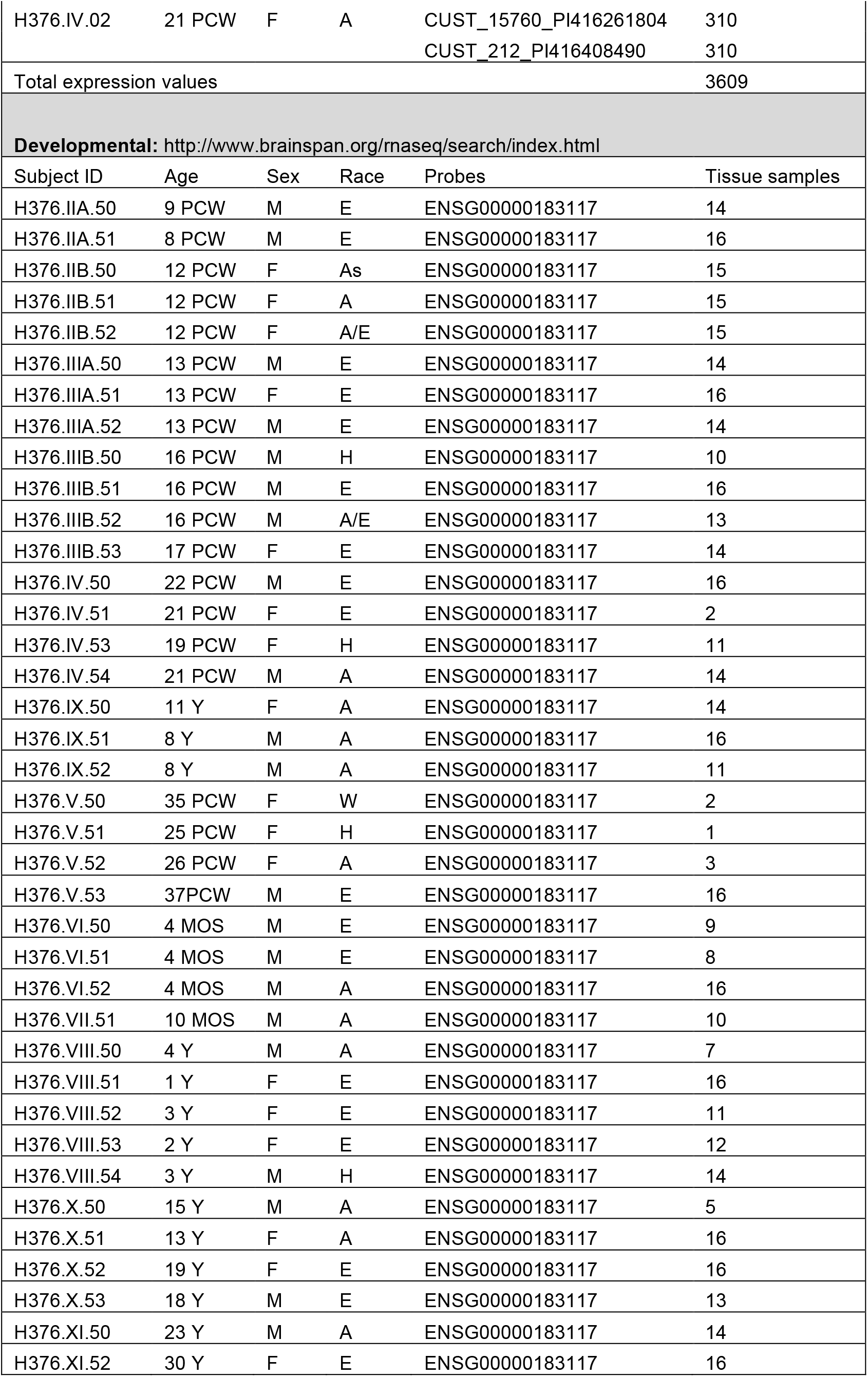
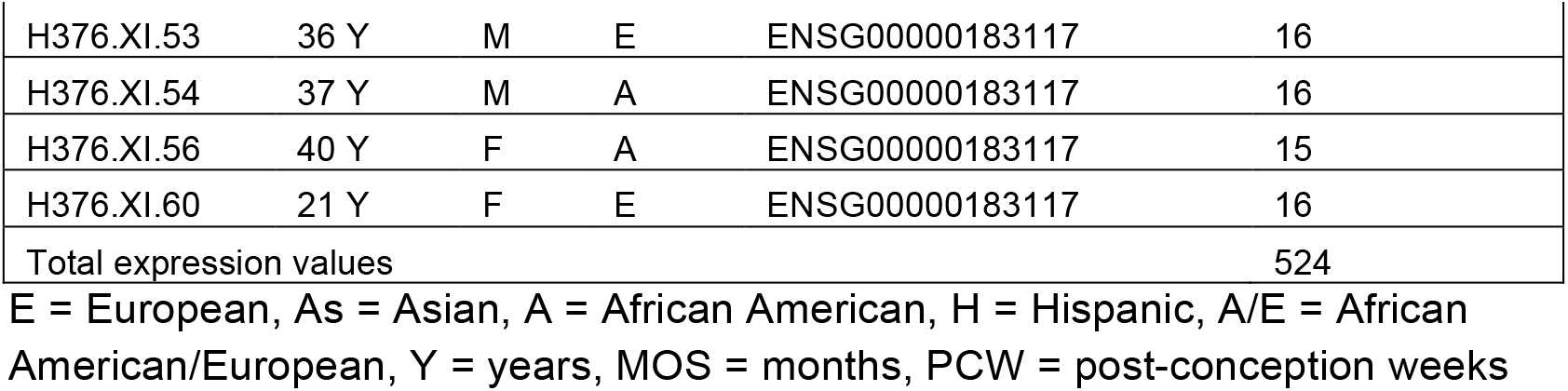
Metadata for samples used in the spatial and temporal analysis of *CSMD1* expression from the Allen Brain Sciences Institute.

To assess the temporal pattern of expression for the whole brain and the whole amygdala, we used the RNA-sequencing dataset of brains across the lifespan from the BrainSpan Atlas of the Developing Human Brain http://www.brainspan.org/). This dataset underwent quality control and normalization procedures as described previously (Sunkin et al., 2013) and in technical white papers available at http://human.brain-map.org/ under the respective atlas’s “Documentation” tab. We downloaded *CSMD1* expression RPKM data at http://www.brainspan.org/, which originates from a single probe, 42 brains, and 524 tissue samples from left and right hemispheres making up 26 brain regions (Table 2). We pooled data across brain regions and into developmental groups including early fetal (8 to 13 weeks post-conception), late fetal (16 to 26 weeks post-conception), childhood (4 months to 8 years old), adolescence (13-21 years old), and adulthood (23-40 years old). We performed an ANOVA with post-hoc *t*-tests to test for differences in expression by developmental stage, accepting *p* < 0.05 as significant.

## Results

### GWAS meta-analysis

As seen in Figure 2, the GWAS meta-analysis of our *a priori* amygdala network phenotypes identified over 200 single nucleotide polymorphisms (SNPs) each with associations to one of the three amygdala networks at the suggestive threshold (*p*<1×10^−5^) and one genome-wide significant association for SNP rs10105357 with the medial amygdala network (beta=0.203, *p*=2.97×10^−8^). The quantile-quantile plots showed no genomic inflation (medial: 1.024; dorsal: 1.011; ventrolateral: 1.009; Supplementary Figure 2a-c). Individual regressions for rs10105357 in IMAGEN and PNC made roughly equal contributions to the meta-analysis (Figure 3a). This SNP localizes to an intergenic region on chromosome 8p23.2, in close proximity to *CSMD1* (CUB And Sushi Multiple Domains 1) and sits atop a peak of other strongly associated SNPs in high linkage disequilibrium (Figure 3b). We next analyzed the functional significance of rs10105357, and for exploratory purposes, the top SNPs associated with the dorsal and ventrolateral amygdala networks (the latter detailed in Supplementary Results and Figures).

**Figure 2.**
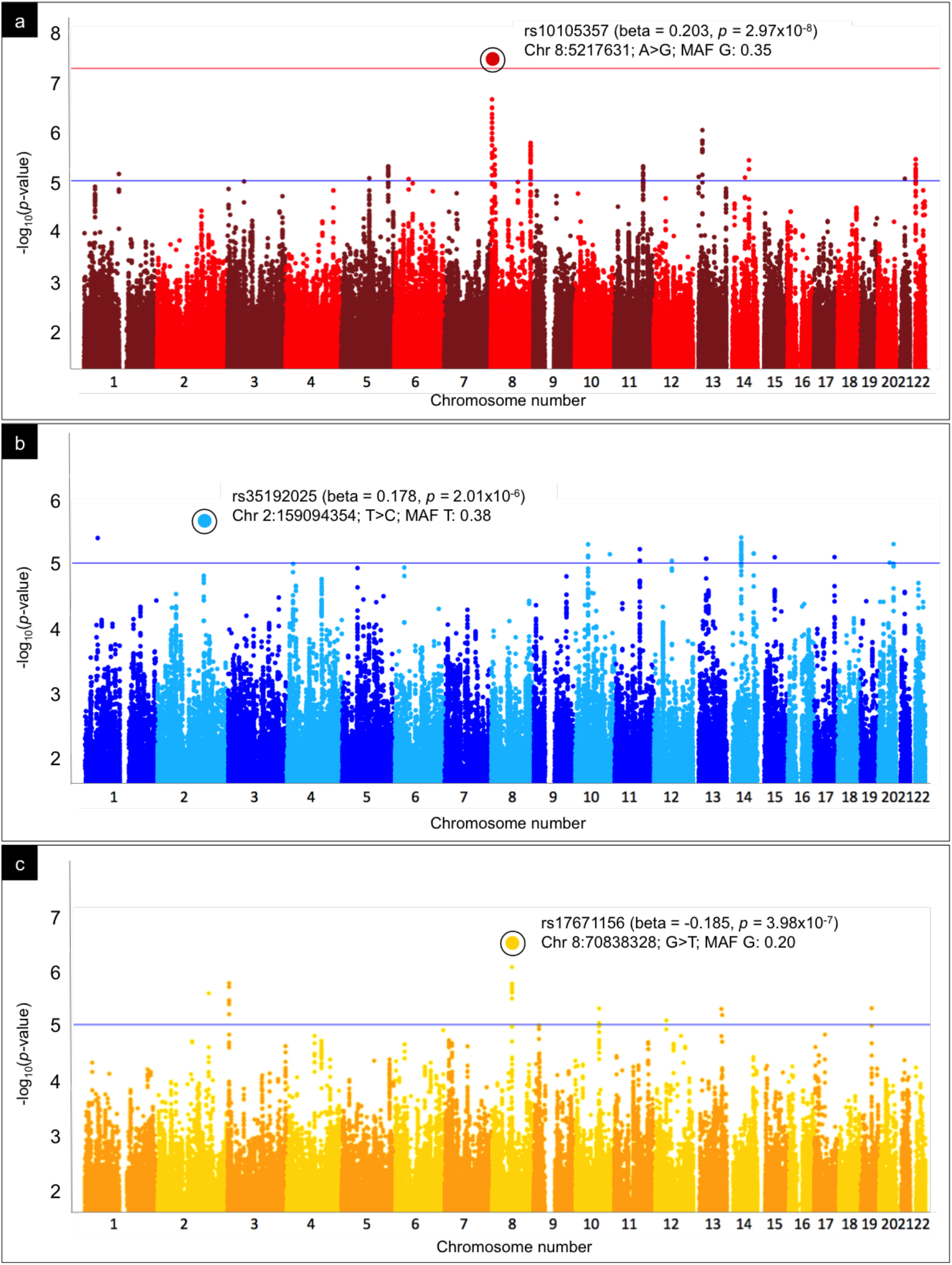
Genetic variants associated with amygdala network phenotypes. Manhattan plots for the (a) medial, (b) dorsal, and (c) ventrolateral amygdala network phenotypes showing SNPs plotted according to chromosomal location (x-axis), with -log10 *p-*values (y-axis) derived from the GWAS meta-analysis. The lower horizontal line indicates the suggestive threshold (*p*<1×10^−5^) and upper horizontal line the threshold for genome-wide statistical significance (*p*<5×10^−8^).

**Figure 3.**
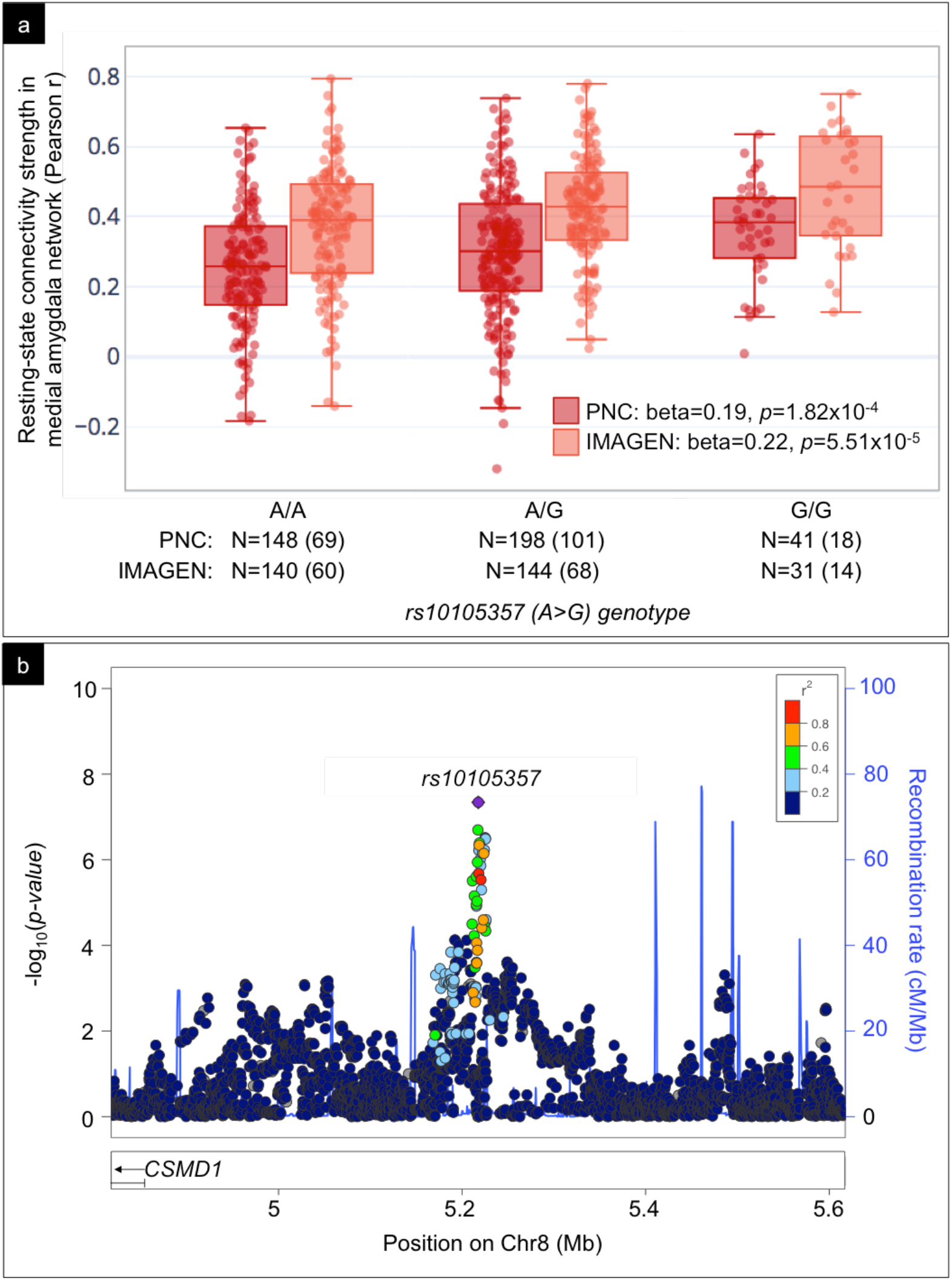
rs10105357 association with medial amygdala network connectivity and locus-centered association plot. (a). Boxplots showing the association of resting-state connectivity strength in the medial amygdala network (y-axis) with rs10105357 for the PNC (dark red) and IMAGEN (light red) cohorts where total number of people (N) with each genotype are listed on the x-axis (number of females in parentheses). Results of the regression for the additive model in each dataset are overlaid after controlling for potential confounds. (b). Regional association plot displaying -log10 *p-*values (y-axis, left) of SNPs as dots against their physical position on the chromosome (x-axis) as well as gene annotations from the UCSC genome browser (below x-axis) and spikes representing estimated recombination rates from the 1000Genomes EUR population (y-axis right) (http://hapmap.ncbi.nlm.nih.gov/). The SNP’s colors indicate LD according to a scale from r^2^ = 0 to r^2^ = 1 (inset in right corner) based on pairwise r^2^ values from 1000Genomes EUR population.

### Functional validation by expression quantitative trait loci (eQTL) analysis

Furthermore, this SNP acts as a significant eQTL for *CSMD1* expression in the temporal cortex (Figure 4a) in the Mayo RNA-Sequencing healthy control dataset (Allen et al., 2016) for the additive model (N=62: beta=0.20, *p*=0.047). Given that the expression pattern suggests a strong recessive effect (Fig 5a), we used an ANOVA and post-hoc *t*-tests for all subsequent validation analyses (N=62: ANOVA F=6.218, *p*=0.004). People with two copies of the minor allele have significantly higher *CSMD1* expression in temporal cortex than people with one or no copy of the allele (t=3.281, *p*=0.002), which remains significant after controlling for age of death, sex, RNA integrity number, and post-mortem intervals (t=2.200, *p*=0.032). Of the two top suggestive SNPs for the other networks, rs17671156-G, which is associated with lower connectivity in the ventrolateral amygdala network, acts as a significant eQTL for temporal cortex, in this case decreasing *SLCO5A1* expression (detailed in Supplementary Results and Supplementary Figure 3).

**Figure 4.**
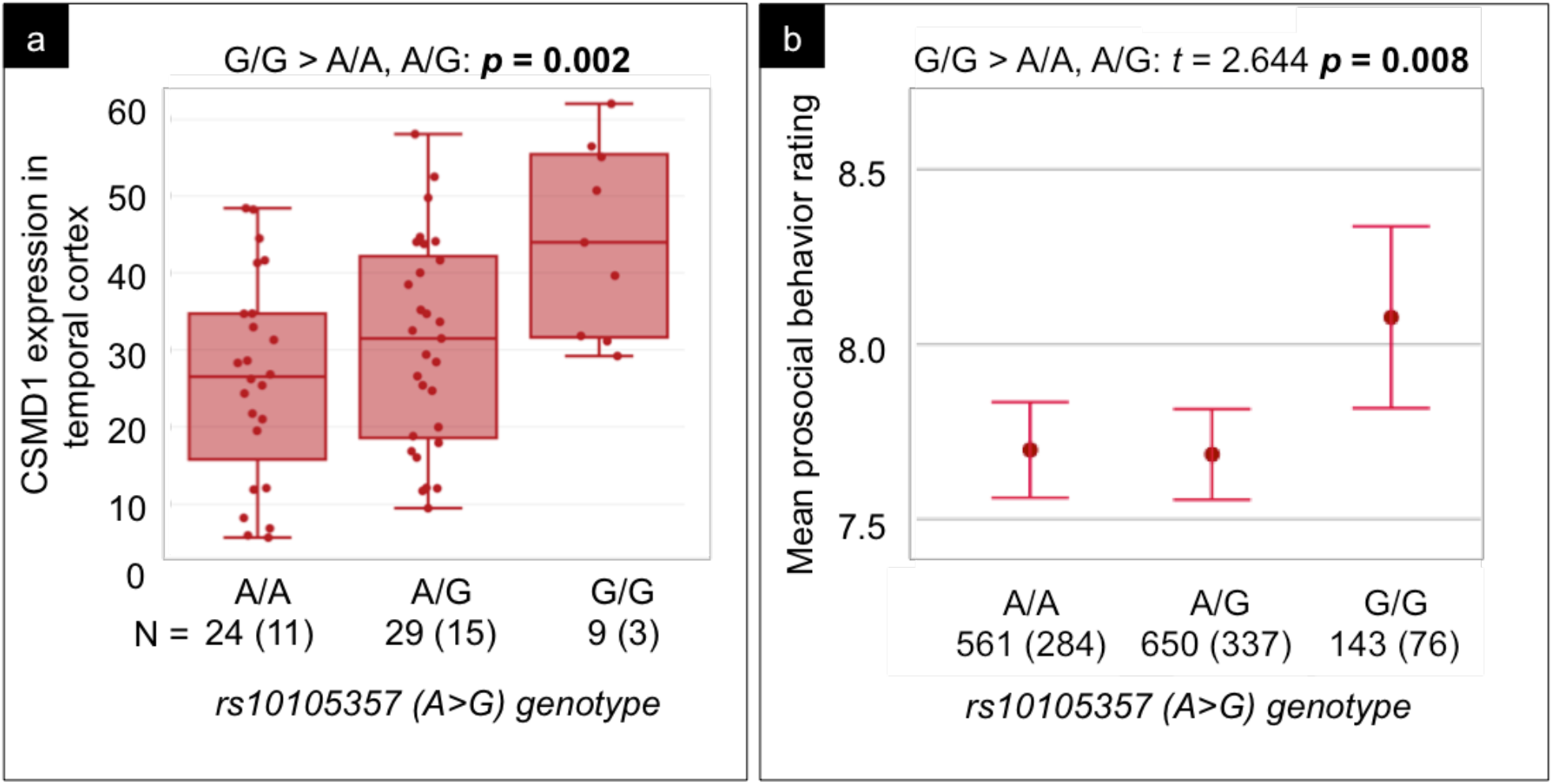
rs10105357 is associated with increased *CSMD1* expression and prosocial behavior. (a). Boxplot showing the effect of rs10105357 on *CSMD1* expression based on eQTL analysis of the Mayo RNA-sequencing control dataset (N=62) where total number of people (N) with each genotype are listed on the x-axis (number of female in parentheses) and *CSMD1* expression on the y-axis (normalized FPKM=Fragments Per Kilobase of transcript per Million mapped reads). Unadjusted *p-*value from the *t-*test for the recessive model is overlaid. (b). An error bar plot showing mean prosocial behavior ratings (y-axis) stratified by genotype (x-axis) with sample characteristics for each genotype. Dots represent means and error bars represent standard error. Results of the *t-*test for the recessive model are overlaid.

**Figure 5.**
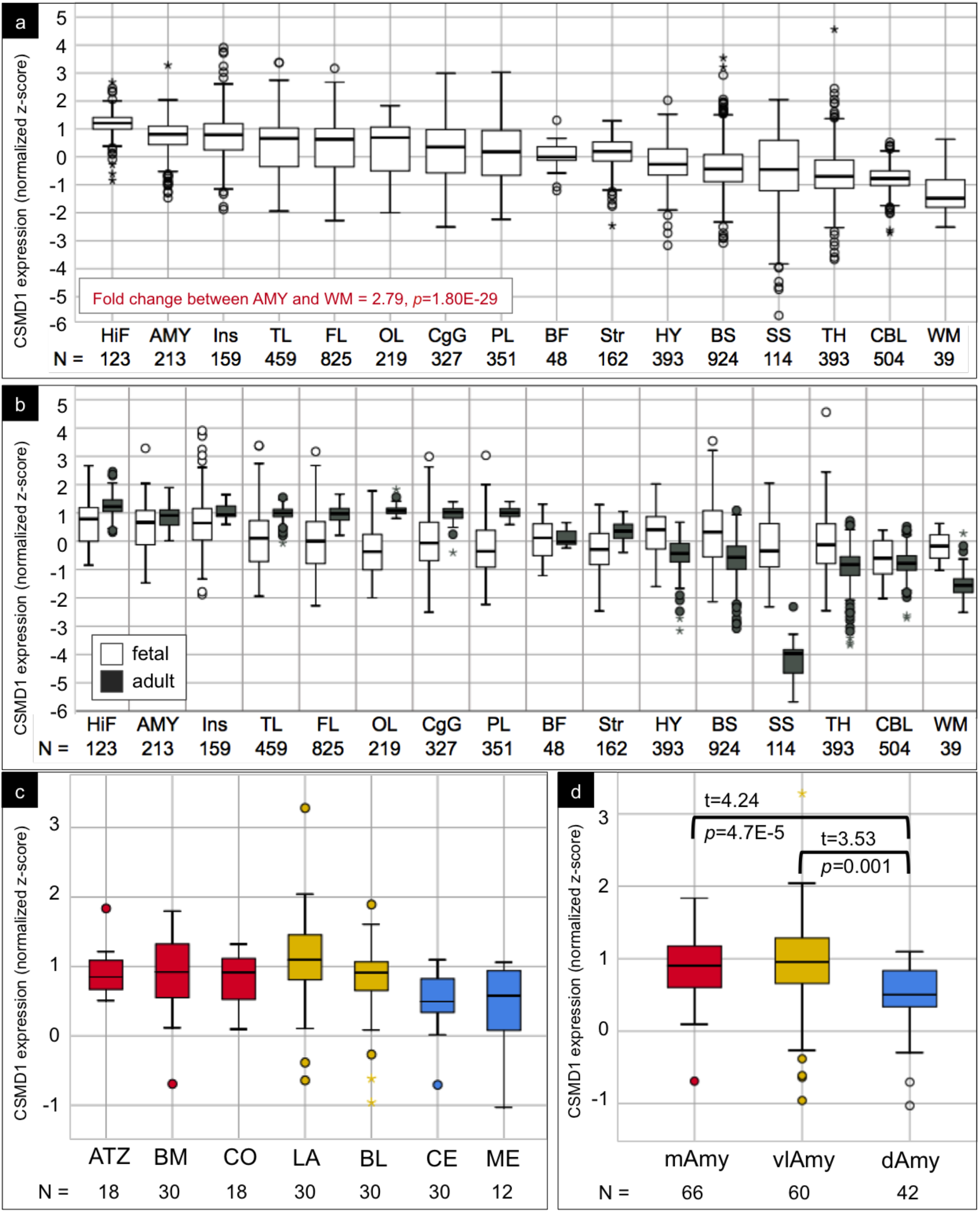
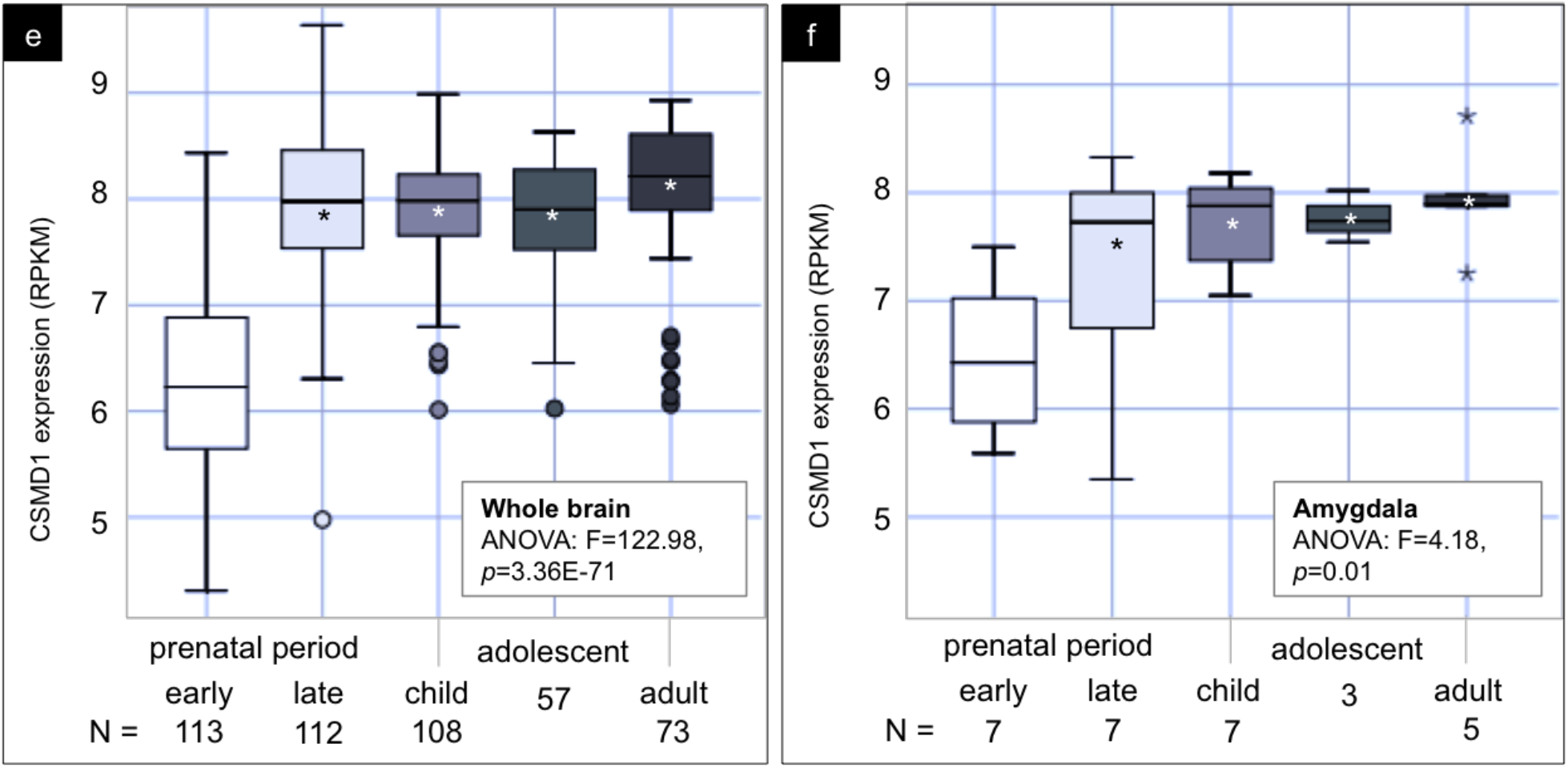
*CSMD1* expression in the brain and amygdala across the lifespan. Boxplots a-d display normalized z-scores for *CSMD1* expression pooled across fetal (http://www.brainspan.org/lcm/) and adult (http://human.brain-map.org/) microarray datasets for (a). all brain regions, (b). all brain regions stratified by dataset, (c). amygdala nuclei color coded by subregions belonging to each amygdala network and (d). the expression pooled across these nuclei for each amygdala subregion. Boxplots e-f display RPKM *CSMD1* expression values from the developmental RNA-sequencing dataset (http://www.brainspan.org/) stratified by age groups for the (e). whole brain without the amygdala and (f). the whole amygdala (* on the in the boxes indicate post-hoc *t*-tests with *p*<0.05, least significant difference corrected, comparing early prenatal group to each subsequent age group). For all boxplots, * above or below the boxes indicate outliers, N=number of expression values pooled into each box. Abbreviations: HiF=hippocampal formation, AMY=amygdala, Ins=insula, TL=temporal lobe, FL=frontal lobe, OL=occipital lobe, CgG=cingulate gyrus, PL=parietal lobe, BF=basal forebrain, Str=striatum, HY=hypothalamus, BS=brainstem, TH=thalamus, SS=somatosensory cortex, CBL=cerebellum, WM=white matter, ATZ=amygdalohippocampal transition zone, BM=basomedial nucleus of the amygdala, CO=cortical nuclei of the amygdala, LA=lateral nucleus of the amygdala, BL=basolateral nucleus of the amygdala, CE=central nucleus of the amygdala, ME=medial nucleus of the amygdala, mAmy=medial amygdala subregion, vlAmy=ventrolateral amygdala subregion, dAmy=dorsal amygdala subregion.

### Functional validation by endophenotype analysis

We next interrogated the endophenotype concept by testing the effect of genotype for our top SNPs on behavioral phenotypes corresponding to each amygdala network in an independent subset of our cohorts that were not included in the GWAS (sample characteristics stratified by genotype detailed in Table 1 and Figure 4b). We found a main effect of genotype for rs10105357 on prosocial behavior (one way ANOVA F=3.50, *p*=0.030), in which adolescents with two copies of the minor allele, rs10105357-G, participate in more prosocial behaviors than adolescents with one or no copy of the allele (t=2.644, *p*=0.008; Figure 4b), which remains significant after controlling for potential confounds (i.e., sex, handedness, Tanner stage of puberty, imaging center, and three genetic principal components; t=2.08, *p*=0.038).

### *CSMD1* expression in the brain and amygdala across the lifespan

We next investigated the localization of *CSMD1* expression and its cross-sectional developmental trajectory using three datasets publicly available from the Allen Brain Sciences Institute (http://human.brain-map.org/). Across the lifespan, ranging from 8 weeks post-conception to 57 years old, we found highest *CSMD1* expression in the cortex overall, and particularly in the hippocampus and amygdala (2.79 fold higher than white matter, *p*=1.80×10^−29^; Figure 5a-b). Nuclei within the broader medial and ventrolateral subregions of the amygdala have significantly higher expression than the dorsal subregion (Figure 5c-d). *CSMD1* expression is greatest later in development for the brain as a whole as well as for the amygdala (Figure 5e-f). The amygdala shows significantly greater expression of *CSMD1* in late prenatal period, childhood, adolescence, and adulthood as compared with the early prenatal period (Figure 5f). Further details regarding data sources, sample demographics, and statistics for this analysis can be found in the Methods.

## Discussion

Here, we show for the first time that large-scale resting-state networks of the amygdala can serve as endophenotypes for social behavior. Prior work has explored the amygdala’s connectional architecture with resting-state fMRI, defining three networks that support distinct aspects of social behavior, including social perception, affiliation, and aversion (Bickart et al., 2012). The present study builds on this to investigate the genetic drivers of these amygdala networks using a GWAS meta-analysis of two independent healthy adolescent samples. We discovered a common genome-wide SNP, rs10105357, associated with variations in connectivity in the amygdala’s medial network. This network emanates from voxels in the vicinity of its cortical and basomedial subnuclei and includes regions important for reward processing and goal-based decisions in general (red network in Figure 1), and affiliative behaviors in the social realm (Bickart, Dickerson, et al., 2014). For example, in humans, regions within this network respond to positive social feedback (e.g. approval) (Izuma, Saito, & Sadato, 2010) that elicit prosocial sentiments (e.g. compassion or empathy) (Tabibnia, Satpute, & Lieberman, 2008; Zahn, de Oliveira-Souza, Bramati, Garrido, & Moll, 2009) and in turn motivate decisions to behave altruistically and cooperate (e.g. donation to charities or repaying trust in kind) (Moll et al., 2006; Rilling, Sanfey, Aronson, Nystrom, & Cohen, 2004). People with stronger resting-state connectivity within this network have larger, more complex social networks (Bickart et al., 2012). Alternatively, people with frontotemporal dementia who have more atrophy in this network show greater impairments in empathy, warmth, and intimacy, and sometimes even behave coldly or cruelly (Bickart, Brickhouse, et al., 2014).

We found that rs10105357 lies in close proximity to *CSMD1* and has a functional role in regulating the expression of this gene in the temporal cortex in an independent sample of adults from the Mayo RNA-Sequencing control dataset (Allen et al., 2016). This is particularly compelling because *CSMD1* is a biologically plausible gene for the regulation of amygdala circuit development and function. *CSMD1* is a large gene that spans more than 2 Mb on the short arm of chromosome 8. *CSMD1* expression occurs predominantly in the brain in rodents and humans (Escudero-Esparza, Kalchishkova, Kurbasic, Jiang, & Blom, 2013). In rodents, expression is particularly extensive in the medial temporal lobe and increases through development reaching a maximum in adulthood (Kraus et al., 2006). Here, we also confirmed this rough spatial and temporal pattern of expression for the first time in the human brain, and in the amygdala in particular, using three datasets publicly available from the Allen Brain Sciences Institute. Leveraging 2929 tissue samples in 52 brains ranging in age from 8 weeks post-conception to 57 years old, we found that *CSMD1* expression is greatest in medial temporal lobe structures including the amygdala (Figure 5a-b), particularly in nuclei of the medial and ventrolateral subregions (Figure 5c-d), with greatest expression in later stages of development (Figure 5e-f). *CSMD1* encodes a large, nearly 400kDa, transmembrane protein that colocalizes with F-actin in the growth cone and filopodia of embryonic neurons (Kraus et al., 2006). The CSMD1 protein is highly conserved across species and like its closely related family members, CSMD2 and CSMD3, is composed of alternating CUB and Sushi domains that share strong homologies with other developmentally-regulated complement-control related proteins (Kraus et al., 2006). *In vitro*, recombinant versions of CSMD1 inhibit the classical complement pathway by blocking conversion and deposition of C3 in rats (Kraus et al., 2006) and promote degradation of C3 and C4 in humans (Escudero-Esparza et al., 2013).

Based on these features, researchers hypothesize that *CSMD1* plays a direct role in refining the precise wiring of developing brain circuits by regulating axonal growth towards targets for synaptogenesis (Kraus et al., 2006) as well as in inhibiting aberrant complement-mediated pruning of synapses (Steen et al., 2013). Consistent with this hypothesis, genetic variants likely related to dysregulated *CSMD1* expression have been linked to neuropsychiatric conditions such as schizophrenia (Kwon, Wang, & Tsai, 2013), autism (Krumm et al., 2015), major depression (Sullivan et al., 2008), and bipolar disorder (Xu et al., 2014). Carriers of such variants show disruption in normal white matter connectivity globally (Giddaluru et al., 2016) and resting-state connectivity in the default mode network (Meda et al., 2014) as well as impairment in several neurocognitive functions (Koiliari et al., 2014). A knockout study shows that decreased *CSMD1* expression results in murine phenotypic homologs of anxiety, agoraphobia, depression, and learned helplessness (Steen et al., 2013). Our finding that rs10105357-G relates to both increased amygdala connectivity and *CSMD1* expression in temporal cortex suggests that this variant may confer a beneficial gain of function, presumably to affiliative behavior, in contrast to the risk variants previously discovered.

Further supporting its beneficial effects, we found that adolescents who carry two minor alleles for rs10105357, which results in the highest *CSMD1* expression levels, participate in significantly more prosocial behaviors in an independent subset of healthy adolescents from the IMAGEN cohort. We derived prosocial behavior from the Strengths and Difficulties Questionnaire (SDQ) (Goodman, 1997), which is an efficient, widely used, and validated screen for both adolescent strengths (i.e., prosocial behavior) and difficulties (i.e., emotional and behavioral). The prosocial behavior subscale assesses resources, indexing adaptive competencies that may promote adjustment and protect against emotional and behavioral difficulties (Silva et al., 2015). This subscale has discriminant validity in healthy adolescents, showing highest sensitivity and specificity for identifying prosocial resources at a cutoff score of 8/10 when compared to a more comprehensive evaluation of social skills from the Development and Well-being Assessment (Silva et al., 2015). It also has clinical utility. For example, lower prosocial behavior scores can be seen in children with autism, mean of 4.5/10 (Christakou et al., 2013), and their siblings, mean of 7.0/10 (Hastings, 2003), as compared with control samples, mean of 8.6/10. In the present study, adolescents carrying two copies of the rs10105357 minor allele scored a mean of 8.1/10 on the prosocial behavior subscale as compared to 7.7/10 for their peers carrying one or no minor alleles (*p*=0.008). Taken together, this finding suggests that being a rs10105357-G homozygote provides some social advantage, promoting social competence and/or protecting against problems with the development of such skills.

This work builds on prior imaging genetic studies of the amygdala by performing a whole genome meta-analysis on a comprehensive model of amygdala circuitry. To date, prior studies have mostly conducted targeted analyses on specific SNPs or genes with measures of amygdala size, functional activity during tasks, and in a few cases, connectivity with a limited set of target regions. These variations have been found to drive differences in amygdala structure (Good et al., 2003; Meyer-Lindenberg et al., 2006b), reactivity (Furmark et al., 2004; Meyer-Lindenberg et al., 2006b; Tost et al., 2010), and connectivity (Buckholtz et al., 2008; Kruschwitz et al., 2015; Mothersill et al., 2014), which have been linked to differences in aspects of sociality, such as social phobia (Furmark et al., 2004), face processing (Mothersill et al., 2014) and social temperament (Buckholtz et al., 2008; Meyer-Lindenberg et al., 2006b; Tost et al., 2010). Fewer studies have conducted GWAS on amygdala imaging phenotypes, including reactivity to faces displaying anger or fear in a mixed populations of healthy people and patients with bipolar disorder and schizophrenia that identified a SNP near *PHOX2B* (Ousdal et al., 2012) and reactivity to hostile-appearing faces in a combined sample of patients with bipolar disorder and healthy controls that identified a SNP near *DOK5* (Liu et al., 2010).

Despite the compelling findings of this study, we acknowledge several shortcomings. First, the samples used for the GWAS were relatively small, containing ∼700 adolescents overall. They were also homogenous in terms of ethnicity, age, and health (with no prior systemic, neurologic, or psychiatric diseases). These features limit generalizability and thus warrant replication in larger samples as well as other ethnic groups, ages, and particularly the vulnerable populations most at risk for the neuropsychiatric diseases presumed to be affected by abnormal function of the genes and disruptions of the networks investigated here.

Overall, this study provides strong support for the endophenotype concept (Meyer-Lindenberg & Weinberger, 2006), suggesting that the amygdala networks may be valuable in reducing the search space and increasing statistical power for the discovery of genetic variations that affect social behavior. Indeed, the top SNPs from our GWAS of the dorsal and ventrolateral amygdala networks also have relevant links to social behavior (described in detail in Supplementary Results and Discussion and Supplementary Figure 3-5). Perhaps most compelling though, the top SNP for the ventrolateral amygdala network, rs17671156, drives decreases in connectivity in that network as well as decreases in *SLCO5A1* expression in temporal lobe tissue in an independent sample of healthy adult brains, and relates to decreased performance in social perception tasks in two independent samples of healthy adolescents. In the clinical realm, amygdala network endophenotyping has the potential to accelerate genetic discovery in disorders of social function, such as autism. Future work will aim to further understand the role of *CSMD1* as both a diagnostic marker and therapeutic target for such conditions. *CSMD1* is a particularly compelling target because it may be accessible in vivo from spinal fluid (Luykx et al., 2014) and is a complement control related protein under investigation as a player in the immune hypothesis of neuropsychiatric disease (Réus et al., 2015).

## Acknowledgments

This work received support from the following sources: The Feldman Foundation CA, the European Union-funded FP6 Integrated Project IMAGEN (Reinforcement-related behaviour in normal brain function and psychopathology) (LSHM-CT-2007-037286), the Horizon 2020 funded ERC Advanced Grant ‘STRATIFY’ (Brain network based stratification of reinforcement-related disorders) (695313), ERANID (Understanding the Interplay between Cultural, Biological and Subjective Factors in Drug Use Pathways) (PR-ST-0416-10004), BRIDGET (JPND: BRain Imaging, cognition Dementia and next generation GEnomics) (MR/N027558/1), the FP7 projects IMAGEMEND (602450; IMAging GEnetics for MENtal Disorders) and MATRICS (603016), the Innovative Medicine Initiative Project EU-AIMS (115300-2), the Medical Research Council Grant ‘c-VEDA’ (Consortium on Vulnerability to Externalizing Disorders and Addictions) (MR/N000390/1), the Swedish Research Council FORMAS, the Medical Research Council, the National Institute for Health Research (NIHR) Biomedical Research Centre at South London and Maudsley NHS Foundation Trust and King’s College London, the Bundesministeriumfür Bildung und Forschung (BMBF grants 01GS08152; 01EV0711; eMED SysAlc01ZX1311A; Forschungsnetz AERIAL 01EE1406A, 01EE1406B), the Deutsche Forschungsgemeinschaft (DFG grants SM 80/7-2, SFB 940/2, NE 1383/14-1), the Medical Research Foundation and Medical research council (grant MR/R00465X/1), the Human Brain Project (HBP SGA 2). Further support was provided by grants from: ANR (project AF12-NEUR0008-01 - WM2NA, and ANR-12-SAMA-0004), the Fondation de France, the Fondation pour la Recherche Médicale, the Mission Interministérielle de Lutte-contre-les-Drogues-et-les-Conduites-Addictives (MILDECA), the Assistance-Publique-Hôpitaux-de-Paris and INSERM (interface grant), Paris Sud University IDEX 2012; the National Institutes of Health, Science Foundation Ireland (16/ERCD/3797), U.S.A. (Axon, Testosterone and Mental Health during Adolescence; RO1 MH085772-01A1), and by NIH Consortium grant U54 EB020403, supported by a cross-NIH alliance that funds Big Data to Knowledge Centres of Excellence, the Medical Research Council (grant number MR/L016311/1).

IMAGEN consortium: Tobias Banaschewski M.D., Ph.D., Gareth J. Barker Ph.D., Arun L.W. Bokde Ph.D., Uli Bromberg Ph.D., Christian Büchel M.D., Erin Burke Quinlan, PhD, Sylvane Desrivières Ph.D., Herta Flor Ph.D., Antoine Grigis Ph.D., Vincent Frouin, Hugh Garavan Ph.D., Penny Gowland Ph.D., Andreas Heinz M.D., Ph.D., Bernd Ittermann Ph.D., Jean-Luc Martinot M.D., Ph.D., Marie-Laure Paillère Martinot M.D., Ph.D., Eric Artiges M.D., Ph.D., Herve Lemaitre Ph.D., Frauke Nees Ph.D., Dimitri Papadopoulos Orfanos Ph.D., Tomáš Paus M.D., Ph.D., Luise Poustka M.D., Sarah Hohmann M.D., Sabina Millenet Dipl.-Psych., Juliane H. Fröhner Dipl.-Psych., Michael N. Smolka M.D., Henrik Walter M.D., Ph.D., Robert Whelan Ph.D., Gunter Schumann M.D.

## Contributions

Acquired the data: Banaschewski, T; Bokde, ALW; Flor, H; Garavan, H; Gowland, P; Heinz, A; Ittermann, B; Martinot, J-L; Paillère Martinot, M-L; Nees, F; Paus, T; Smolka, Newsom, M; Contributed unpublished reagents/analytic tools: Quinlan, EB; Desrivières, S; Papadopoulos Orfanos, D; Poustka, L; Fröhner, JH; Walter, H; Schumann, G; Design study: Bickart, KC; Napolioni, V; Sadaghiani, S; Greicius, MD; Neuroimaging processing: Ng, B Khan, RR; Newsom, M; Richiardi, J; Altmann, A; Artiges, E; Whelan, R; Neuroimaging processing: Ng, B; Khan, RR; Newsom, M; Richiardi, J; Altmann, A; Connectivity analysis: Bickart, KC, Khan, RR; Newsom, M Genetic analysis: Napolioni, V; Khan, RR; Richiardi, J; Altmann, A; Expression quantitative trait loci analysis: Kim, Y; Khan, RR; Behavioral analysis: Bickart, KC, Sadaghiani, S; *CSMD1* expression analysis: Bickart, KC, Richiardi, J; Writing and editing: Bickart, KC, Napolioni, V; Khan, RR; Kim, Y; Richiardi, J; Altmann, A; Greicius, MD

## References

1000 Genomes Project Consortium, Auton, A., Brooks, L. D., Durbin, R. M., Garrison, E. P., Kang, H. M., … Abecasis, G. R. (2015). A global reference for human genetic variation. Nature, 526(7571), 68–74.

Allen, M., Carrasquillo, M. M., Funk, C., Heavner, B. D., Zou, F., Younkin, C. S., … Ertekin-Taner, N. (2016). Human whole genome genotype and transcriptome data for Alzheimer’s and other neurodegenerative diseases. Scientific Data, 3, 160089.

Behzadi, Y., Restom, K., Liau, J., & Liu, T. T. (2007). A component based noise correction method (CompCor) for BOLD and perfusion based fMRI. NeuroImage, 37(1), 90–101.

Bickart, K. C., Brickhouse, M., Negreira, A., Sapolsky, D., Barrett, L. F., & Dickerson, B. C. (2014). Atrophy in distinct corticolimbic networks in frontotemporal dementia relates to social impairments measured using the Social Impairment Rating Scale. Journal of Neurology, Neurosurgery, and Psychiatry, 85(4), 438–448.

Bickart, K. C., Dickerson, B. C., & Barrett, L. F. (2014). The amygdala as a hub in brain networks that support social life. Neuropsychologia, 63, 235–248.

Bickart, K. C., Hollenbeck, M. C., Barrett, L. F., & Dickerson, B. C. (2012). Intrinsic Amygdala-Cortical Functional Connectivity Predicts Social Network Size in Humans. Journal of Neuroscience, 32(42), 14729–14741.

Buckholtz, J. W., Callicott, J. H., Kolachana, B., Hariri, A. R., Goldberg, T. E., Genderson, M., … Meyer-Lindenberg, A. (2007). Genetic variation in MAOA modulates ventromedial prefrontal circuitry mediating individual differences in human personality. Molecular Psychiatry, 13(3), 313–324.

Buckholtz, J. W., Callicott, J. H., Kolachana, B., Hariri, A. R., Goldberg, T. E., Genderson, M., … Meyer-Lindenberg, A. (2008). Genetic variation in MAOA modulates ventromedial prefrontal circuitry mediating individual differences in human personality. Molecular Psychiatry, 13(3), 313–324.

Canli, T., & Lesch, K.-P. (2007). Long story short: the serotonin transporter in emotion regulation and social cognition. Nature Neuroscience, 10(9), 1103–1109.

Chang, C. C., Chow, C. C., Tellier, L. C., Vattikuti, S., Purcell, S. M., & Lee, J. J. (2015). Second-generation PLINK: rising to the challenge of larger and richer datasets. GigaScience, 4, 7.

Christakou, A., Murphy, C. M., Chantiluke, K., Cubillo, A. I., Smith, A. B., Giampietro, V., … Rubia, K. (2013). Disorder-specific functional abnormalities during sustained attention in youth with Attention Deficit Hyperactivity Disorder (ADHD) and with autism. Molecular Psychiatry, 18(2), 236–244.

Delaneau, O., Marchini, J., 1000 Genomes Project Consortium, & 1000 Genomes Project Consortium. (2014). Integrating sequence and array data to create an improved 1000 Genomes Project haplotype reference panel. Nature Communications, 5, 3934.

Escudero-Esparza, A., Kalchishkova, N., Kurbasic, E., Jiang, W. G., & Blom, A. M. (2013). The novel complement inhibitor human CUB and Sushi multiple domains 1 (CSMD1) protein promotes factor I-mediated degradation of C4b and C3b and inhibits the membrane attack complex assembly. FASEB Journal: Official Publication of the Federation of American Societies for Experimental Biology, 27(12), 5083–5093.

Fakhoury, M. (2018). Imaging genetics in autism spectrum disorders: Linking genetics and brain imaging in the pursuit of the underlying neurobiological mechanisms. Progress in Neuro-Psychopharmacology & Biological Psychiatry, 80(Pt B), 101–114.

Freese, J. L., & Amaral, D. G. (2009). Neuroanatomy of the primate amygdala. In P. J. Whalen & E. A. Phelps (Eds.), The human amygdala (pp. 3–42). New York, NY, US: Guilford Press.

Furmark, T., Tillfors, M., Garpenstrand, H., Marteinsdottir, I., Långström, B., Oreland, L., & Fredrikson, M. (2004). Serotonin transporter polymorphism related to amygdala excitability and symptom severity in patients with social phobia. Neuroscience Letters, 362(3), 189–192.

Giddaluru, S., Espeseth, T., Salami, A., Westlye, L. T., Lundquist, A., Christoforou, A., … Nyberg, L. (2016). Genetics of structural connectivity and information processing in the brain. Brain Structure & Function, 221(9), 4643–4661.

Good, C. D., Lawrence, K., Thomas, N. S., Price, C. J., Ashburner, J., Friston, K. J., … Skuse, D. H. (2003). Dosage sensitive X linked locus influences the development of amygdala and orbitofrontal cortex, and fear recognition in humans. Brain: A Journal of Neurology, 126(11), 2431–2446.

Goodman, R. (1997). The Strengths and Difficulties Questionnaire: a research note. Journal of Child Psychology and Psychiatry, and Allied Disciplines, 38(5), 581–586.

Gur, R. E., Moore, T. M., Calkins, M. E., Ruparel, K., & Gur, R. C. (2017). Face Processing Measures of Social Cognition: A Dimensional Approach to Developmental Psychopathology. Biological Psychiatry: Cognitive Neuroscience and Neuroimaging. http://doi.org/10.1016/j.bpsc.2017.03.010

Hastings, R. P. (2003). Brief report: Behavioral adjustment of siblings of children with autism. Journal of Autism and Developmental Disorders, 33(1), 99–104.

Hawrylycz, M. J., Lein, E. S., Guillozet-Bongaarts, A. L., Shen, E. H., Ng, L., Miller, J. A., … Jones, A. R. (2012). An anatomically comprehensive atlas of the adult human brain transcriptome. Nature, 489(7416), 391–399.

Howie, B. N., Donnelly, P., & Marchini, J. (2009). A Flexible and Accurate Genotype Imputation Method for the Next Generation of Genome-Wide Association Studies. PLoS Genetics, 5(6), e1000529.

Izuma, K., Saito, D. N., & Sadato, N. (2010). Processing of the incentive for social approval in the ventral striatum during charitable donation. Journal of Cognitive Neuroscience, 22(4), 621–631.

Kent, W. J. (2002). The Human Genome Browser at UCSC. Genome Research, 12(6), 996–1006.

Koiliari, E., Roussos, P., Pasparakis, E., Lencz, T., Malhotra, A., Siever, L. J., … Bitsios, P. (2014). The CSMD1 genome-wide associated schizophrenia risk variant rs10503253 affects general cognitive ability and executive function in healthy males. Schizophrenia Research, 154(1-3), 42–47.

Kraus, D. M., Elliott, G. S., Chute, H., Horan, T., Pfenninger, K. H., Sanford, S. D., … Holers, V. M. (2006). CSMD1 is a novel multiple domain complement-regulatory protein highly expressed in the central nervous system and epithelial tissues. Journal of Immunology, 176(7), 4419–4430.

Krumm, N., Turner, T. N., Baker, C., Vives, L., Mohajeri, K., Witherspoon, K., … Eichler, E. E. (2015). Excess of rare, inherited truncating mutations in autism. Nature Genetics, 47(6), 582–588.

Kruschwitz, J. D., Walter, M., Varikuti, D., Jensen, J., Plichta, M. M., Haddad, L., … Walter, H. (2015). 5-HTTLPR/rs25531 polymorphism and neuroticism are linked by resting state functional connectivity of amygdala and fusiform gyrus. Brain Structure & Function, 220(4), 2373–2385.

Kwon, E., Wang, W., & Tsai, L.-H. (2013). Validation of schizophrenia-associated genes CSMD1, C10orf26, CACNA1C and TCF4 as miR-137 targets. Molecular Psychiatry, 18(1), 11–12.

Li, P., Fan, T.-T., Zhao, R.-J., Han, Y., Shi, L., Sun, H.-Q., … Lu, L. (2017). Altered Brain Network Connectivity as a Potential Endophenotype of Schizophrenia. Scientific Reports, 7(1), 5483.

Liu, X., Akula, N., Skup, M., Brotman, M. A., Leibenluft, E., & McMahon, F. J. (2010). A genome-wide association study of amygdala activation in youths with and without bipolar disorder. Journal of the American Academy of Child and Adolescent Psychiatry, 49(1), 33–41.

Luykx, J. J., Bakker, S. C., Lentjes, E., Neeleman, M., Strengman, E., Mentink, L., … Ophoff, R. A. (2014). Genome-wide association study of monoamine metabolite levels in human cerebrospinal fluid. Molecular Psychiatry, 19(2), 228–234.

Mägi, R., & Morris, A. P. (2010). GWAMA: software for genome-wide association meta-analysis. BMC Bioinformatics, 11, 288.

Meda, S. A., Ruaño, G., Windemuth, A., O’Neil, K., Berwise, C., Dunn, S. M., … Pearlson, G. D. (2014). Multivariate analysis reveals genetic associations of the resting default mode network in psychotic bipolar disorder and schizophrenia. Proceedings of the National Academy of Sciences of the United States of America, 111(19), E2066–75.

Meyer-Lindenberg, A., Buckholtz, J. W., Kolachana, B., R Hariri, A., Pezawas, L., Blasi, G., … Weinberger, D. R. (2006a). Neural mechanisms of genetic risk for impulsivity and violence in humans. Proceedings of the National Academy of Sciences of the United States of America, 103(16), 6269–6274.

Meyer-Lindenberg, A., Buckholtz, J. W., Kolachana, B., R Hariri, A., Pezawas, L., Blasi, G., … Weinberger, D. R. (2006b). Neural mechanisms of genetic risk for impulsivity and violence in humans. Proceedings of the National Academy of Sciences of the United States of America, 103(16), 6269–6274.

Meyer-Lindenberg, A., & Weinberger, D. R. (2006). Intermediate phenotypes and genetic mechanisms of psychiatric disorders. Nature Reviews. Neuroscience, 7(10), 818–827.

Miller, J. A., Ding, S.-L., Sunkin, S. M., Smith, K. A., Ng, L., Szafer, A., … Lein, E. S. (2014). Transcriptional landscape of the prenatal human brain. Nature, 508(7495), 199–206.

Moll, J., Krueger, F., Zahn, R., Pardini, M., de Oliveira-Souza, R., & Grafman, J. (2006). Human fronto-mesolimbic networks guide decisions about charitable donation. Proceedings of the National Academy of Sciences of the United States of America, 103(42), 15623–15628.

Mothersill, O., Morris, D. W., Kelly, S., Rose, E. J., Fahey, C., O’Brien, C., … Donohoe, G. (2014). Effects of MIR137 on fronto-amygdala functional connectivity. NeuroImage, 90, 189–195.

Nacewicz, B. M., Dalton, K. M., Johnstone, T., Long, M. T., McAuliff, E. M., Oakes, T. R., … Davidson, R. J. (2006). Amygdala volume and nonverbal social impairment in adolescent and adult males with autism. Archives of General Psychiatry, 63(12), 1417–1428.

Nusbaum, C., Mikkelsen, T. S., Zody, M. C., Asakawa, S., Taudien, S., Garber, M., … Lander, E. S. (2006). DNA sequence and analysis of human chromosome 8. Nature, 439(7074), 331–335.

Olweus, D. (1996). Bully/victim problems in school. Prospects, 26(2), 331–359.

Ousdal, O. T., Anand Brown, A., Jensen, J., Nakstad, P. H., Melle, I., Agartz, I., … Andreassen, O. A. (2012). Associations between variants near a monoaminergic pathways gene (PHOX2B) and amygdala reactivity: a genome-wide functional imaging study. Twin Research and Human Genetics: The Official Journal of the International Society for Twin Studies, 15(3), 273–285.

Price, A. L., Patterson, N. J., Plenge, R. M., Weinblatt, M. E., Shadick, N. A., & Reich, D. (2006). Principal components analysis corrects for stratification in genome-wide association studies. Nature Genetics, 38(8), 904–909.

Réus, G. Z., Fries, G. R., Stertz, L., Badawy, M., Passos, I. C., Barichello, T., … Quevedo, J. (2015). The role of inflammation and microglial activation in the pathophysiology of psychiatric disorders. Neuroscience, 300, 141–154.

Rilling, J. K., Sanfey, A. G., Aronson, J. A., Nystrom, L. E., & Cohen, J. D. (2004). Opposing BOLD responses to reciprocated and unreciprocated altruism in putative reward pathways. Neuroreport, 15(16), 2539–2543.

Rosen, H. J., Perry, R. J., Murphy, J., Kramer, J. H., Mychack, P., Schuff, N., … Miller, B. L. (2002). Emotion comprehension in the temporal variant of frontotemporal dementia. Brain: A Journal of Neurology, 125(10), 2286–2295.

Satterthwaite, T. D., Elliott, M. A., Ruparel, K., Loughead, J., Prabhakaran, K., Calkins, M. E., … Gur, R. E. (2014). Neuroimaging of the Philadelphia neurodevelopmental cohort. NeuroImage, 86, 544–553.

Schumann, C. M., Bauman, M. D., & Amaral, D. G. (2011). Abnormal structure or function of the amygdala is a common component of neurodevelopmental disorders. Neuropsychologia, 49(4), 745–759.

Schumann, G., Loth, E., Banaschewski, T., Barbot, A., Barker, G., Büchel, C., … Struve, M. (2010). The IMAGEN study: reinforcement-related behaviour in normal brain function and psychopathology. Molecular Psychiatry, 15(12), 1128–1139.

Silva, T. B. F., Osório, F. L., & Loureiro, S. R. (2015). SDQ: discriminative validity and diagnostic potential. Frontiers in Psychology, 6, 811.

Steen, V. M., Nepal, C., Ersland, K. M., Holdhus, R., Nævdal, M., Ratvik, S. M., … Håvik, B. (2013). Neuropsychological deficits in mice depleted of the schizophrenia susceptibility gene CSMD1. PloS One, 8(11), e79501.

Sullivan, P. F., de Geus, E. J. C., Willemsen, G., James, M. R., Smit, J. H., Zandbelt, T., … Penninx, B. W. J. H. (2008). Genome-wide association for major depressive disorder: a possible role for the presynaptic protein piccolo. Molecular Psychiatry, 14, 359.

SNPs associated with CSMD1 were found to be in top significant SNPs to associate with MDD, as found in supplemental excel table.

Sunkin, S. M., Ng, L., Lau, C., Dolbeare, T., Gilbert, T. L., Thompson, C. L., … Dang, C. (2013). Allen Brain Atlas: an integrated spatio-temporal portal for exploring the central nervous system. Nucleic Acids Research, 41(Database issue), D996–D1008.

Tabibnia, G., Satpute, A. B., & Lieberman, M. D. (2008). The sunny side of fairness: preference for fairness activates reward circuitry (and disregarding unfairness activates self-control circuitry). Psychological Science, 19(4), 339–347.

Tost, H., Kolachana, B., Hakimi, S., Lemaitre, H., Verchinski, B. A., Mattay, V. S., … Meyer-Lindenberg, A. (2010). A common allele in the oxytocin receptor gene (OXTR) impacts prosocial temperament and human hypothalamic-limbic structure and function. Proceedings of the National Academy of Sciences of the United States of America, 107(31), 13936–13941.

Xu, W., Cohen-Woods, S., Chen, Q., Noor, A., Knight, J., Hosang, G., … Vincent, J. B. (2014). Genome-wide association study of bipolar disorder in Canadian and UK populations corroborates disease loci including SYNE1 and CSMD1. BMC Medical Genetics, 15, 2.

Yang, Y., Raine, A., Narr, K. L., Colletti, P., & Toga, A. W. (2009). Localization of deformations within the amygdala in individuals with psychopathy. Archives of General Psychiatry, 66(9), 986–994.

Zahn, R., de Oliveira-Souza, R., Bramati, I., Garrido, G., & Moll, J. (2009). Subgenual cingulate activity reflects individual differences in empathic concern. Neuroscience Letters, 457(2), 107–110.

