## Supplemental material for "Genetic variation in *CSMD1* affects amygdala connectivity and prosocial behavior"

### Supplementary Information

#### Methods

Given the function of the dorsal amygdala network, we hypothesized that carriers of the rs35192025-C polymorphism would demonstrate phenotypic differences in instruments assessing tendencies to judge and avoid threatening people. The IMAGEN cohort underwent a relevant assessment, specifically, the Bully/Victim Questionnaire [15]. It assesses prevalence of participating in bullying behaviors and being the victim of such behaviors. Items included, 'A student/ peer left me out of things on purpose, excluded me from their group of friends or completely ignored me.', 'I was hit, kicked, pushed or shoved around, or locked indoors by a student/ peer.', 'I took part in bullying another student/ peer at school.', 'I called another student/ peer mean names, made fun of, or teased him or her in a hurtful way.', 'I kept a student/ peer out of things on purpose, excluded that student/peer from my group of friends, or completely ignored that student/ peer.', 'I hit, kicked, pushed, shoved around, or locked a student/ peer indoors.', 'I have been bullied by a teacher.', 'I have been bullied by a family member.'. Respondents rate each item on a 5 point scale (1 'None' 2 'Only once or twice' 3 '2 or 3 times a month' 4 'About once a week' 5 'Several times a week'). We computed a “bully to victim” ratio where a score greater than 1 indicates the respondent endorsed a higher prevalence of bullying than being bullied, and the reverse for scores less than 1.

Given the function of the ventrolateral amygdala network, we hypothesized that carriers of the rs17671156-T polymorphism would demonstrate phenotypic differences in instruments testing aspects of social perception. The PNC and IMAGEN cohorts underwent face processing tasks. The PNC tasks included face memory, facial expression identification, facial expression intensity discrimination, and face age determination. Based on a prior study using the PNC data [16], we z-transformed and averaged the data across the four face processing tasks to produce scores for response time for correct responses. IMAGEN subjects completed a single face processing task of facial expression identification from 4 continua of morphed faces between pairs of expressions. The task generates event and summary scores. Event scores included probability of choosing one emotion or the other in the pair on each continua for each intensity level (e.g., 0, 10, 20, etc) and response time to make this choice, whether correct or incorrect. For summary scores, a threshold value interpolates the intensity point at which subjects began identifying the second facial expression of the pair more than 50% of the time, an estimate of sensitivity or bias for one of the two emotions in each pair. The other summary variable is a noise score, which characterizes the degree of certainty or precision for discriminating facial expressions at the extreme ends of the morph spectrum (e.g., how often subject selects the anger over sad facial expression in faces showing 100% anger or 100% sad; higher values suggests more uncertainty). We chose to assess this value as the other values did not assess response time for correct responses as in the PNC tasks, but rather response time for all events and bias towards one emotion or another.

### Results

#### GWAS

For exploratory purposes, we investigated the top SNPs associated with the dorsal and ventrolateral amygdala networks (connectivity stratified by genotype for these SNPs in PNC and IMAGEN datasets can be seen in Supplementary Figure 1).

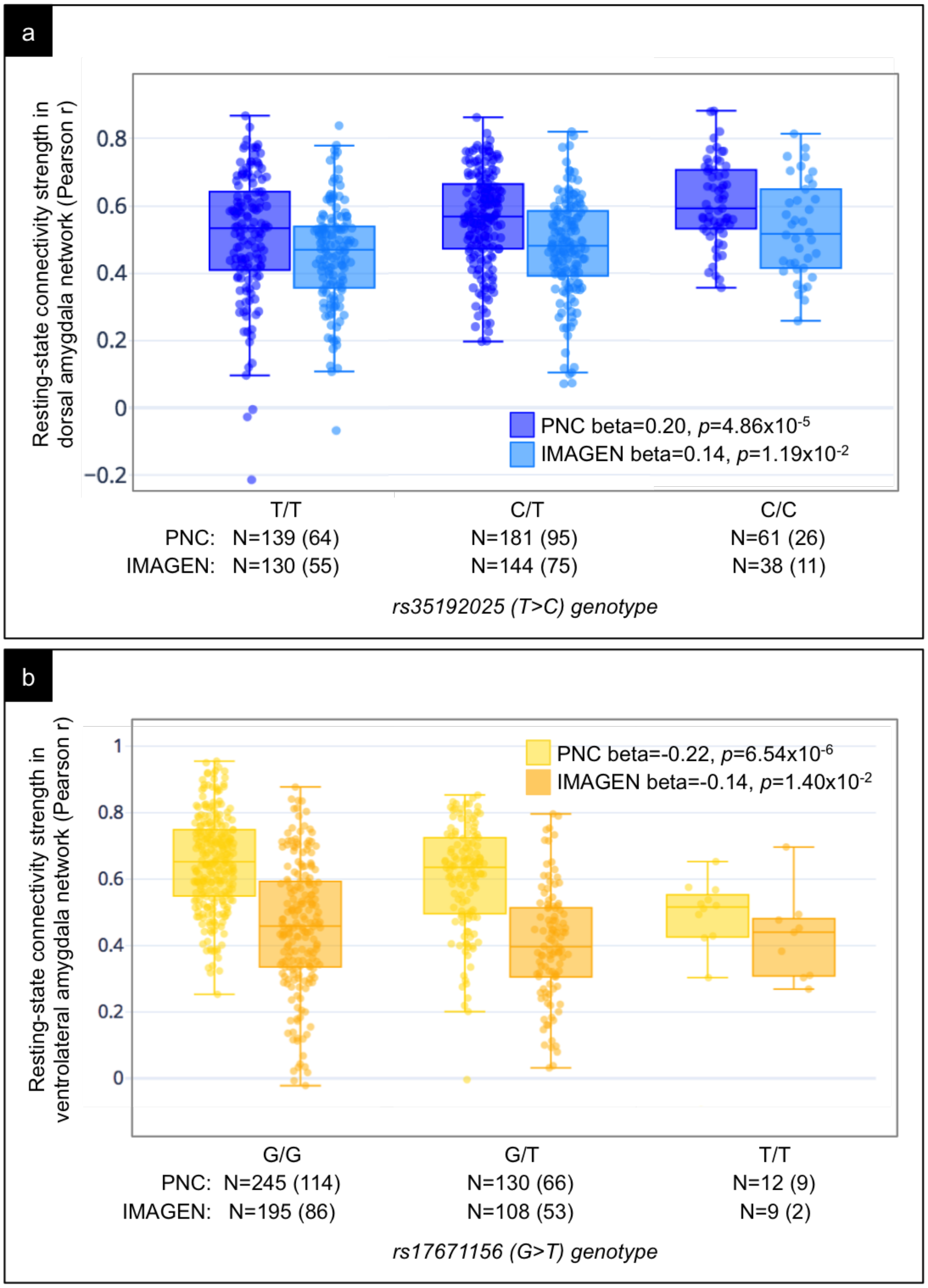


**Supplementary Figure 1 Amygdala network phenotypes stratified by top SNP genotypes and datasets.** Boxplots showing the association of resting-state connectivity strength in the medial amygdala network (y-axis) with rs10105357 for the PNC (dark red) and IMAGEN (light red) datasets where total number of people (N) with each genotype are listed on the x-axis (number of females in parentheses). Results of the regression for the additive model in each dataset are overlaid after controlling for potential confounds.

The quantile-quantile plots for the three GWAS analyses showed no genomic inflation (medial: 1.024; dorsal:1.011; ventrolateral: 1.009; Supplementary Figure 2a-c).

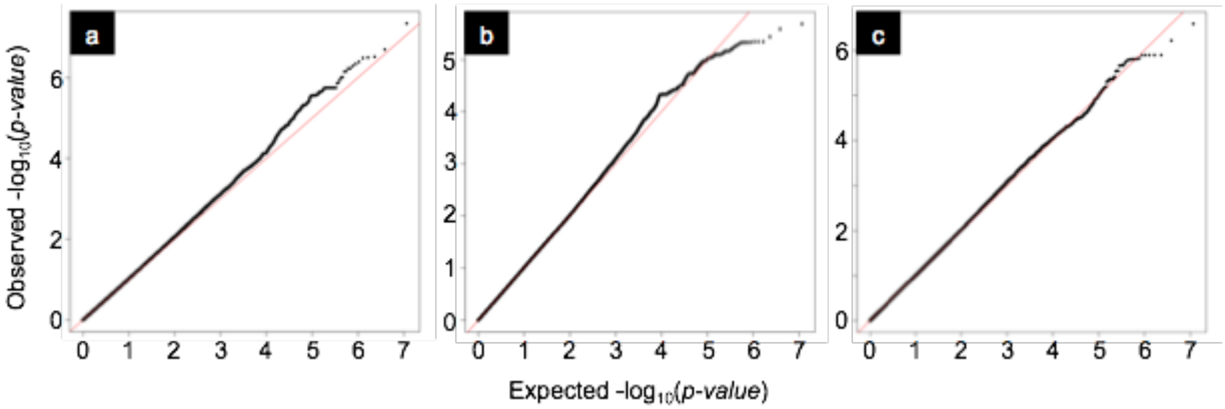


**Supplementary Figure 2 Quantile-quantile plots**. Quantile-quantile plots for the GWAS analyses of medial amygdala network (a), dorsal amygdala network (b), and ventrolateral amygdala network (c).

#### LD and expression quantitative trait loci (eQTL) analysis for top suggestive SNPs:

The top SNPs for the dorsal and ventrolateral amygdala networks also sit atop peaks of strongly associated, highly linked SNPs (see Supplementary Figure 3a and c). Rs35192025 is located within the *CCDC148* gene, on chromosome 2q24.1 (Supplementary Figure 3a). It does not act as a significant eQTL for *CCDC148* expression in the temporal cortex (Supplementary Figure 3b) in the Mayo RNA-Sequencing control dataset [1] (N=62: linear regression beta=-0.034, $p$=0.752 ; ANOVA F=1.574, $p$=0.216). Rs17671156 is located on chromosome 8q13.3, approximately 100Kb upstream from the *SLCO5A1* gene (Supplementary Figure 3c). It acts as a significant eQTL for *SLCO5A1* expression in the temporal cortex (Supplementary Figure 3d), decreasing its expression, in the Mayo RNA-Sequencing control dataset [1] (N=62: linear regression beta=-0.211, $p$=0.063; ANOVA F=7.014, $p$=0.002). People with one or two copies of the minor allele have significantly less *SLCO5A1* expression than people with no copy of the allele (t=-3.672, $p$=0.001), which remains significant after controlling for age of death, sex, RNA integrity number, and post-mortem intervals (t=-3.756, $p$<0.001).

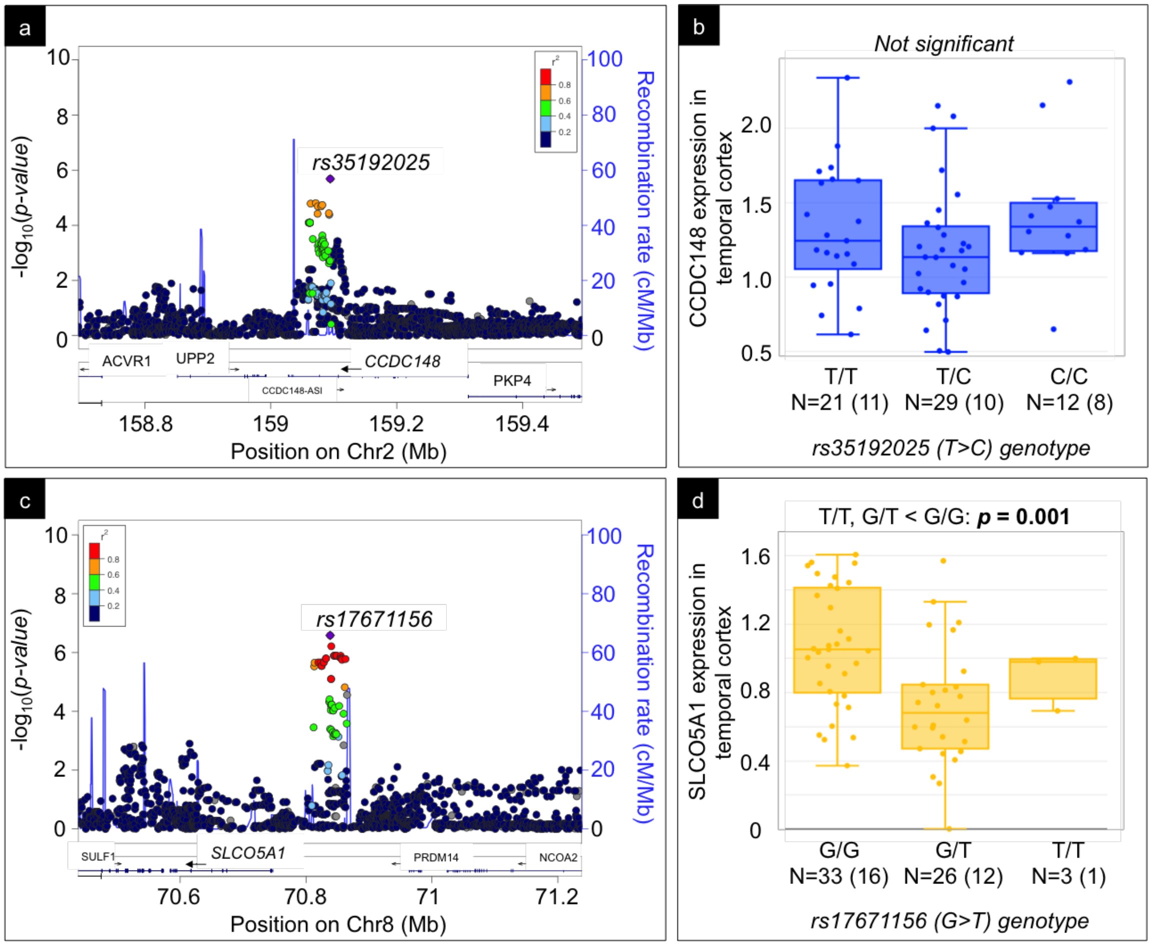


**Supplementary Figure 3 Regional association and eQTL for rs35192025 and rs17671156.** (a, c). Regional association plots displaying -log10 *p*-values (y axis left) of SNPs as dots against their physical position on the chromosome (x-axis) as well as gene annotations from the UCSC genome browser (below x axis) and spikes representing estimated recombination rates from the 1000Genomes EUR population (y-axis right). The SNP's colors indicate LD according to a scale from $r^2$ = 0 to $r^2$ = 1 (inset in right corner) based on pairwise $r^2$ values from 1000Genomes EUR population. (b, d). Boxplot showing no significant effect of rs35192025 on *CCDC148* (b) and a significant effect of rs17171156 on *SLCO5A1* (d) expression in temporal cortex based on eQTL analysis of the Mayo RNA-Sequencing control dataset (N=62) where total number of people with each genotype are listed on the x axis (female in parentheses) and gene expression on the y-axis (normalized FPKM). Unadjusted *p*-value from the *t*-test for the dominant model overlaid for rs17671156.

#### Endophenotype analysis for top suggestive SNPs:

In the IMAGEN cohort, there is a main effect of rs35192025 on the bully to victim ratio (One way ANOVA F= 3.619, p=0.027). In post hoc comparisons, adolescents with one or two copies of the minor allele show a significantly lower bully to victim ratio than their peers with no copy of the minor allele (t=-2.674, p=0.008; Figure 4), which remains significant after controlling for potential confounds (t=2.79, p=0.005).

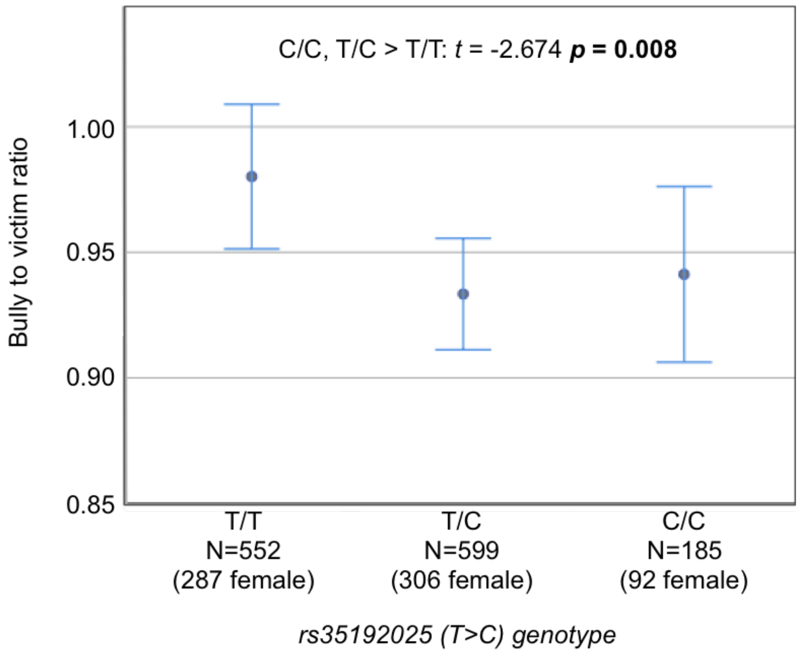


**Supplementary Figure 4 Error bar plot for bully to victim ratio by rs35192025 genotype.** An error bar plot showing mean bully to victim ratings (y axis) stratified by genotype (x axis) with sample characteristics for each genotype. Results of the *t*-test for the dominant model overlaid.

In the PNC cohort, there is a main effect of rs17671156 on social perception across four face processing tasks (Response time: F=3.80, *p*=0.023), in which adolescents with two copies of the minor allele show significantly slower response times for correct trials than their peers with one or no copy of the minor allele (Response time: t=2.72, *p*=0.007; Figure 5a), which remains significant after controlling for potential confounds (t=2.73, *p*=0.007). In the IMAGEN cohort, there is a trend for a main effect of rs17671156 on face processing uncertainty in the emotion identification from morphed faces task (One way ANOVA F=3.36, p=0.065). In post hoc comparisons, adolescents with one or two copies of the minor allele show significantly greater uncertainty on the task as compared to their peers with no copy of the minor allele (t=2.181, p=0.029; Figure 5b), which remains significant after controlling for potential confounds (t=2.458, p=0.014).

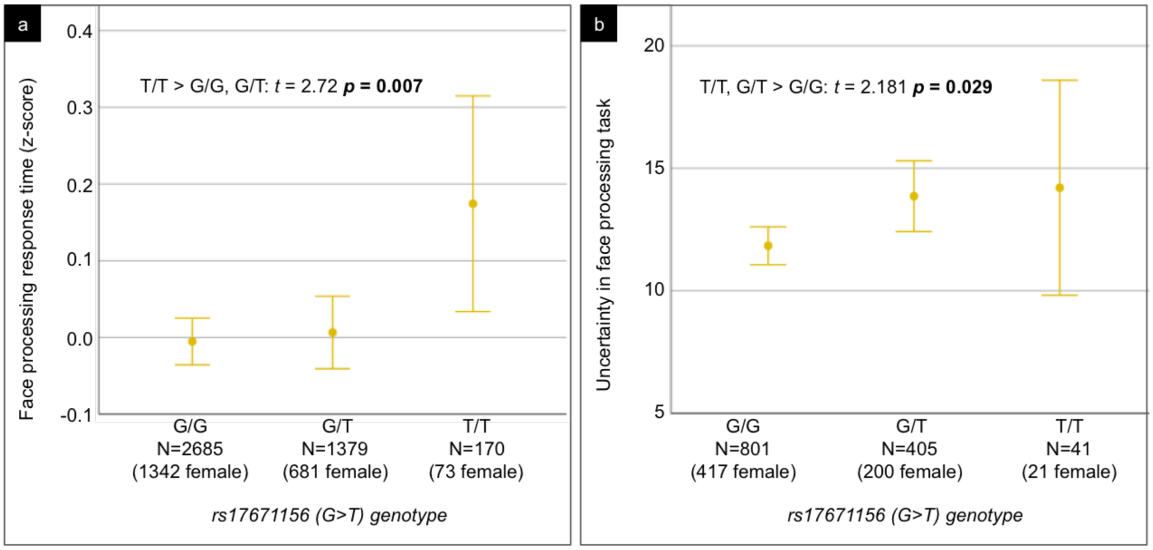


**Supplementary Figure 5 Error bar plot for face processing response time and uncertainty by rs17671156 genotype.** Error bar plots showing (a) response times z-scores for correct responses on 4 face processing tasks (y-axis) in the PNC cohort stratified by genotype (x-axis) with sample characteristics for each genotype. Results of the *t*-test for the recessive model overlaid. An error bar plot showing (b) uncertainty scores on an emotion identification face processing task (y-axis) in the IMAGEN cohort stratified by genotype (x-axis) with sample characteristics for each genotype. Results of the *t*-test for the dominant model overlaid.

#### Supplementary Discussion:

For exploratory purposes, we also tested the endophenotype hypothesis on the top suggestive SNPs from our GWAS of the other two amygdala networks, finding that they also have relevant links to social behavior.

The top SNP for the ventrolateral amygdala network, rs17671156, drives decreases in connectivity in that network as well as decreases in *SLCO5A1* expression in temporal lobe tissue and decreases in performance in social perception tasks in two independent samples of healthy adolescents. The ventrolateral amygdala network contains brain regions along the ventral visual stream from primary visual cortex through the fusiform gyrus and sensory association areas in other parts of the temporal lobe, including the superior temporal sulcus, as well as the orbitofrontal cortex. In the social realm, these areas are implicated in learning about and interpreting social signals [2]. Electrophysiological recording studies in monkeys demonstrate neurons in the amygdala, lateral orbitofrontal cortex, and temporal lobe visual areas respond to facial identity [3–7] and changeable aspects of faces like expressions, eye gaze, and lip movement [6–9], which all serve as signals for social communication. Task-based fMRI studies in humans demonstrate a similar topography of brain responses to socially salient features in the human face such as facial expressions [10–14], eye gaze [15–17], facial identity [18–22], racial or group identity [23–27], and trustworthiness [28–33]. People with increased connectivity in the ventrolateral amygdala network have larger, more complex social networks [34]. People with frontotemporal dementia who have the greatest atrophy in this network, tend to make less frequent eye contact, have difficulty following and interpreting body language, and are less sensitive to others' facial expressions [2].

*SLCO5A1* (Solute carrier organic anion transporter family, member 5A1) is located on the long arm of chromosome 8 (8q13.3). It is a member of the SLCO super family, expressed highest in fetal and adult brain and heart [35,36], and encodes a transmembrane protein that transports organic solutes, both endogenous and commonly used drugs, across the membrane [37].

The top SNP for the dorsal amygdala network, rs35192025, drives increases in connectivity in that network and decreases in bully as compared to victim behaviors, but does not act as a significant eQTL in temporal lobe tissue. In the IMAGEN sample, adolescents with one or two minor alleles are less likely to bully others than to be bullied. Bullying was assessed using the Bully/Victim Questionnaire [38], which is a widely used validated instrument for assessing the prevalence and risks for bullying behaviors and victims of bullying. The dorsal amygdala network contains nociceptive regions responsive to painful, threatening, and biologically important stimuli, including many components of the salience network [39], such as the caudal anterior cingulate, anterior insula, somatosensory operculum, lateral striatum, hypothalamus, thalamus, and several brainstem structures [2]. In the social realm, regions within this network are responsive to untrustworthy-appearing faces and negative social feedback (e.g. disapproval or violations of trust) [28–30,32,40–44] that elicit sentiments of social aversion (e.g. disgust or contempt) [45–50] and in turn motivate decisions to defect cooperation or disengage from a group [51,52].

### References


1. Allen, M. *et al.* Human whole genome genotype and transcriptome data for Alzheimer's and other neurodegenerative diseases. *Sci Data* **3**, 160089 (2016).
2. Bickart, K. C., Dickerson, B. C. & Barrett, L. F. The amygdala as a hub in brain networks that support social life. *Neuropsychologia* **63**, 235–248 (2014).
3. Rolls, E. T. Neurons in the cortex of the temporal lobe and in the amygdala of the monkey with responses selective for faces. *Hum. Neurobiol.* **3**, 209–222 (1984).
4. Leonard, C. M., Rolls, E. T., Wilson, F. A. & Baylis, G. C. Neurons in the amygdala of the monkey with responses selective for faces. *Behav. Brain Res.* **15**, 159–176 (1985).
5. Rolls, E. T., Critchley, H. D., Browning, A. S. & Inoue, K. Face-selective and auditory neurons in the primate orbitofrontal cortex. *Exp. Brain Res.* **170**, 74–87 (2006).
6. Rolls, E. T. The representation of information about faces in the temporal and frontal lobes. *Neuropsychologia* **45**, 124–143 (2007).
7. Hasselmo, M. E., Rolls, E. T. & Baylis, G. C. The role of expression and identity in the face-selective responses of neurons in the temporal visual cortex of the monkey. *Behav. Brain Res.* **32**, 203–218 (1989).
8. Haxby, J. V., Hoffman, E. A. & Gobbini, M. I. Human neural systems for face recognition and social communication. *Biol. Psychiatry* **51**, 59–67 (2002).
9. Baylis, G. C., Rolls, E. T. & Leonard, C. M. Selectivity between Faces in the Responses of a Population of Neurons in the Cortex in the Superior Temporal Sulcus of the Monkey. *Brain Res.* **342**, 91–102 (1985).
10. Morris, J. S. *et al.* A differential neural response in the human amygdala to fearful

and happy facial expressions. *Nature* **383**, 812–815 (1996).

11. Allison, T., Puce, A. & McCarthy, G. Social perception from visual cues: role of the STS region. *Trends Cogn. Sci.* **4**, 267–278 (2000).
12. Phillips, M. L. *et al.* A specific neural substrate for perceiving facial expressions of disgust. *Nature* **389**, 495–498 (1997).
13. Haxby, J. V., Hoffman, E. A. & Gobbini, M. I. The distributed human neural system for face perception. *Trends Cogn. Sci.* **4**, 223–233 (2000).
14. Winston, J. S., Vuilleumier, P. & Dolan, R. J. Effects of low-spatial frequency components of fearful faces on fusiform cortex activity. *Curr. Biol.* **13**, 1824–1829 (2003).
15. Kawashima, R. *et al.* A PET study. *Brain* **122**, 779–783 (1999).
16. George, N., Driver, J. & Dolan, R. J. Seen gaze-direction modulates fusiform activity and its coupling with other brain areas during face processing. *Neuroimage* **13**, 1102–1112 (2001).
17. Richeson, J. A., Todd, A. R., Trawalter, S. & Baird, A. A. Eye-Gaze Direction Modulates Race-Related Amygdala Activity. *Group Process. Intergroup Relat.* **11**, 233–246 (2008).
18. Iidaka, T. *et al.* Dissociable neural responses in the hippocampus to the retrieval of facial identity and emotion: an event-related fMRI study. *Hippocampus* **13**, 429–436 (2003).
19. Schwartz, C. E., Wright, C. I., Shin, L. M., Kagan, J. & Rauch, S. L. Inhibited and uninhibited infants ‘grown up’: adult amygdalar response to novelty. *Science* **300**, 1952–1953 (2003).
20. Pourtois, G., Schwartz, S., Seghier, M. L., Lazeyras, F. & Vuilleumier, P. View-independent coding of face identity in frontal and temporal cortices is modulated by familiarity: an event-related fMRI study. *Neuroimage* **24**, 1214–1224 (2005).

21. Gobbini, M. I. & Haxby, J. V. Neural systems for recognition of familiar faces. *Neuropsychologia* **45**, 32–41 (2007).

22. Wright, P. & Liu, Y. Neutral faces activate the amygdala during identity matching. *Neuroimage* **29**, 628–636 (2006).

23. Hart, A. J. *et al.* Differential response in the human amygdala to racial outgroup vs ingroup face stimuli. *Neuroreport* **11**, 2351–2355 (2000).

24. Phelps, E. A. Faces and races in the brain. *Nat. Neurosci.* **4**, 775–776 (2001).

25. Phelps, E. A. *et al.* Performance on indirect measures of race evaluation predicts amygdala activation. *J. Cogn. Neurosci.* **12**, 729–738 (2000).

26. Cunningham, W. A. *et al.* Separable neural components in the processing of black and white faces. *Psychol. Sci.* **15**, 806–813 (2004).

27. Freeman, J. B., Rule, N. O., Adams, R. B., Jr & Ambady, N. The Neural Basis of Categorical Face Perception: Graded Representations of Face Gender in Fusiform and Orbitofrontal Cortices. *Cereb. Cortex* **20**, 1314–1322 (2010).

28. Winston, J. S., Strange, B. A., O'Doherty, J. & Dolan, R. J. Automatic and intentional brain responses during evaluation of trustworthiness of faces. *Nat. Neurosci.* **5**, 277–283 (2002).

29. Engell, A. D., Haxby, J. V. & Todorov, A. Implicit trustworthiness decisions: automatic coding of face properties in the human amygdala. *J. Cogn. Neurosci.* **19**, 1508–1519 (2007).

30. Todorov, A. & Engell, A. D. The role of the amygdala in implicit evaluation of emotionally neutral faces. *Soc. Cogn. Affect. Neurosci.* **3**, 303–312 (2008).

31. Bzdok, D. *et al.* ALE meta-analysis on facial judgments of trustworthiness and attractiveness. *Brain Struct. Funct.* **215**, 209–223 (2011).

32. Todorov, A., Baron, S. G. & Oosterhof, N. N. Evaluating face trustworthiness: a model based approach. *Soc. Cogn. Affect. Neurosci.* **3**, 119–127 (2008).

33. Said, C. P., Baron, S. G. & Todorov, A. Nonlinear amygdala response to face trustworthiness: contributions of high and low spatial frequency information. *J. Cogn. Neurosci.* **21**, 519–528 (2009).

34. Bickart, K. C., Hollenbeck, M. C., Barrett, L. F. & Dickerson, B. C. Intrinsic Amygdala–Cortical Functional Connectivity Predicts Social Network Size in Humans. *J. Neurosci.* **32**, 14729–14741 (2012).

35. Okabe, M. *et al.* Profiling SLCO and SLC22 genes in the NCI-60 cancer cell lines to identify drug uptake transporters. *Mol. Cancer Ther.* **7**, 3081–3091 (2008).

36. Isidor, B. *et al.* Mesomelia-synostoses syndrome results from deletion of SULF1 and SLCO5A1 genes at 8q13. *Am. J. Hum. Genet.* **87**, 95–100 (2010).

37. Hagenbuch, B. & Meier, P. J. The superfamily of organic anion transporting polypeptides. *Biochim. Biophys. Acta* **1609**, 1–18 (2003).

38. Olweus, D. Bully/victim problems in school. *Prospects* **26**, 331–359 (1996).

39. Seeley, W. W. *et al.* Dissociable intrinsic connectivity networks for salience processing and executive control. *Journal of Neuroscience* **27**, 2349–2356 (2007).

40. Willis, M. L., Palermo, R., Burke, D., McGrillen, K. & Miller, L. Orbitofrontal cortex lesions result in abnormal social judgements to emotional faces. *Neuropsychologia* **48**, 2182–2187 (2010).

41. Sanfey, A. G. Social decision-making: insights from game theory and neuroscience. *Science* **318**, 598–602 (2007).

42. Eisenberger, N. I., Lieberman, M. D. & Williams, K. D. Does rejection hurt? An FMRI study of social exclusion. *Science* **302**, 290–292 (2003).

43. Rilling, J. K. *et al.* The neural correlates of the affective response to unreciprocated cooperation. *Neuropsychologia* **46**, 1256–1266 (2008).

44. Klucharev, V., Hytönen, K., Rijpkema, M., Smidts, A. & Fernández, G. Reinforcement learning signal predicts social conformity. *Neuron* **61**, 140–151

(2009).

45. Phillips, M. L. *et al.* Neural responses to facial and vocal expressions of fear and disgust. *Proc. Biol. Sci.* **265**, 1809–1817 (1998).

46. Decety, J., Jackson, P. L., Sommerville, J. A., Chaminade, T. & Meltzoff, A. N. The neural bases of cooperation and competition: an fMRI investigation. *Neuroimage* **23**, 744–751 (2004).

47. Zahn, R. *et al.* The Neural Basis of Human Social Values: Evidence from Functional MRI. *Cereb. Cortex* **19**, 276–283 (2009).

48. Moll, J. *et al.* The moral affiliations of disgust: a functional MRI study. *Cogn. Behav. Neurol.* **18**, 68–78 (2005).

49. Moll, J. *et al.* The self as a moral agent: linking the neural bases of social agency and moral sensitivity. *Soc. Neurosci.* **2**, 336–352 (2007).

50. Buckholtz, J. W. *et al.* The neural correlates of third-party punishment. *Neuron* **60**, 930–940 (2008).

51. Moll, J. *et al.* Human fronto-mesolimbic networks guide decisions about charitable donation. *Proc. Natl. Acad. Sci. U. S. A.* **103**, 15623–15628 (2006).

52. Sanfey, A. G., Rilling, J. K., Aronson, J. A., Nystrom, L. E. & Cohen, J. D. The neural basis of economic decision-making in the Ultimatum Game. *Science* **300**, 1755–1758 (2003).
